# *Candida albicans* drives colorectal cancer progression by inducing hypoxia signaling

**DOI:** 10.1101/2024.12.25.630341

**Authors:** Wanqiu Wang, Mengqi Yang, Fanglei Gong, Zhenyu Zhang, Yanping Ma, Haihuang Li, Yu Zhao, Changzheng Du, Ningning Li, Guiwei He, Kun Sun

**Author notes:** Corresponding authors (W.W.), (G.H.), and (K.S.).

## Abstract

Colorectal cancer (CRC) is the third leading cause of cancer-related mortality worldwide. Gut microbiota, including fungal species, are increasingly implicated in CRC progression, while the molecular mechanisms underlying host-fungal interactions in this context remain poorly understood. Here, we show that *Candida albicans* (*C. albicans*) activates pro-metastatic signaling pathways, MAPK and NF-κB, leading to upregulation of oncogenic transcription factors c-Jun and c-Myc. This cascade stabilizes and activates hypoxia-inducible factor 1α (HIF-1α), a central regulator of tumor metabolic reprogramming and angiogenesis, ultimately fostering a pro-tumorigenic microenvironment. Mechanistically, we identify candidalysin, a secreted peptide toxin of *C. albicans*, as a critical effector that engages epidermal growth factor receptor (EGFR) and toll-like receptor 2 (TLR2) with context-dependent effects across different epithelial models. These findings are further supported by patient-derived colonic organoids. Collectively, our study delineates a *C. albicans*-EGFR/TLR2-ERK/NF-κB-HIF-1α axis that promotes hypoxia-like responses in CRC, revealing a previously underappreciated role of fungal pathogens in shaping the tumor microenvironment and highlighting avenues for future in *vivo* investigation.

## Introduction

Colorectal cancer (CRC) is one of the leading causes of cancer-related mortality worldwide^1^. The majority of CRC deaths are attributed to disease progression and metastasis^2^; therefore, understanding the intricate mechanisms underlying CRC progression is imperative for improving patient outcomes. Numerous factors have been proposed to be involved in CRC metastasis, including the gut microbiota, a diverse community of microorganisms (including bacteria, viruses, and fungi) closely linked to human health and cancer^3–8^. In particular, dysbiosis of the fungal microbiota is implicated in various cancer types^9–14^. For instance, the absence of receptor proteins responsible for recognizing fungi has been linked to the promotion of colorectal tumorigenesis^15,16^. Among the known fungi, *Candida albicans* (*C. albicans*), an opportunistic pathogenic fungus commonly found in the human intestinal tract, can proliferate under conditions of immunosuppression or microbiome dysbiosis and leads to candidiasis^17^. Once adhering to host cells via adhesins, *C. albicans* transitions from its yeast form to a hyphal form, enabling it to penetrate host tissues and invade epithelial barriers^18–20^. Moreover, *C. albicans* secretes virulence factors, such as candidalysin, which can directly damage host cells and activate host immune responses^21^. In CRC, *C. albicans* is more frequently detected in tumors compared to non-cancerous tissues, indicating a potential role in tumorigenesis and progression^22,23^. Recent studies have uncovered important interplay between *C. albicans* and the tumor microenvironment^23–25,26^. In particular, Zhu and colleagues showed that in mouse models, *C. albicans* colonization induces higher tumor load and more severe colitis^26^. However, involvement of *C. albicans* in CRC progression and the molecular mechanism are still largely elusive.

Previous studies have shown that *Candida albicans* can activate MAPK and NF-κB signaling and has been implicated in modulating hypoxia responses in gut^27,28^, however, its roles in the interaction with CRC cells remain to be fully elucidated. In this study, we employed cancer cells and CRC-derived organoids to investigate the cellular responses elicited by *C. albicans* and to determine whether its virulence factor, candidalysin, contributes to the stabilization of HIF-1α and the ensuing pro-tumorigenic events in CRC.

## Results

### C. albicans is associated with CRC progression

To explore the potential association between fungal presence and colorectal cancer progression, we analyzed tumor samples from four CRC patients (two in stage II and other two stage in III). Fungal signals were detected in both stage III samples but not in stage II samples (Figure 1a). Moreover, presence of fungi surrounding tumor cells was also observed (Figure 1b). While limited by sample size, these observations suggest a possible association between fungal presence and advanced disease stage. To validate this result, we investigated its presence and abundance in colorectal adenocarcinoma samples from The Cancer Genome Atlas (TCGA) project^29^. As shown in Figure 1c-d, the abundance of *C. albicans* is significantly increased in late-stage CRC patients and correlates with poorer prognosis. To elucidates the influence of *C. albicans* on CRC, we co-cultured the CRC cell line HCT116 with *C. albicans.* The results showed that *C. albicans* significantly enhanced cell migration (Figure 1e-f). Under hypoxia, this effect persisted even in the presence of colon bacterium *Bacteroides fragilis*. We then examined epithelial-mesenchymal transition (EMT), a crucial process in cancer progression and metastasis. Analysis of EMT marker gene expression confirmed that *C. albicans* infection reduced E-cadherin levels while increasing Vimentin and Slug expression (Figure 1g, Suppl. Figure S1a). Immunofluorescence staining revealed a reduction in E-cadherin at infection sites, with co-localization of E-cadherin signals and the *C. albicans* cell wall, suggesting a direct interaction between *C. albicans* and the HCT116 cell membrane (Figure 1h, Suppl. Figure S1b). Collectively, these findings suggest that *C. albicans* may be associated with features of late-stage CRC progression by promoting cell migration and EMT.

**Figure 1.**
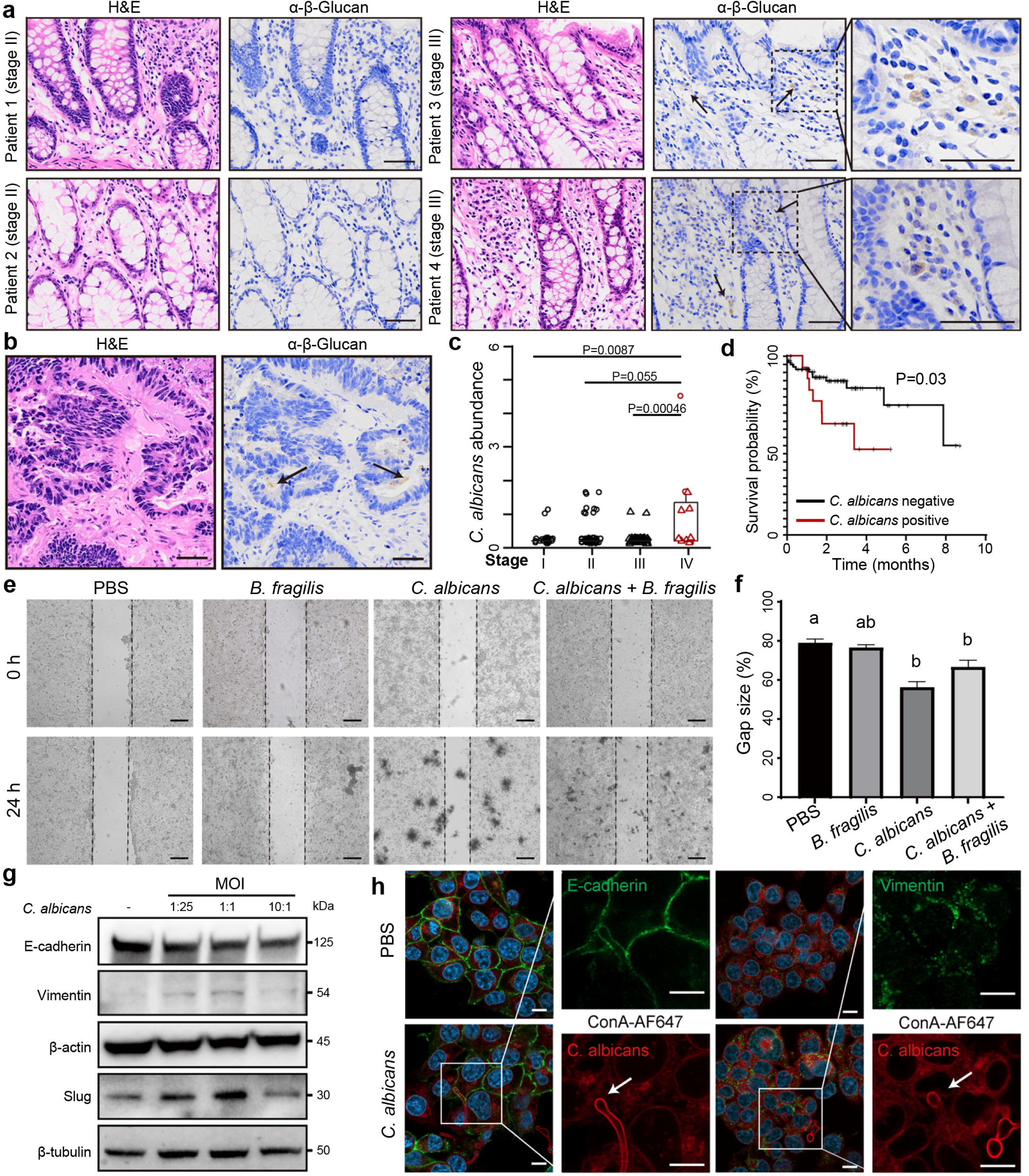
*C. albicans* contributes to late-stage CRC progression. **(a,b)** Representative histological images of tumor tissues from 4 patients stained by hematoxylin and eosin (H&E), antibodies against β-glucan. Samples shown in **b** are derived from the stage III CRC patients. Arrowheads represented positive areas for β-glucan. Scale bars indicated 50 μm. **(c)** Abundance of *C. albicans* in different stages of colorectal adenocarcinoma samples in TCGA project. P-values were calculated using Mann-Whitney U tests. **(d)** Survival curve of colorectal adenocarcinoma patients in TCGA project with or without *C. albicans* infection. P-value was calculated using Log-rank test. **(e, f)** Wound-healing assay for migration capability of PBS-, *B. fragilis*-, *C. albicans-*, and *B. fragilis* with *C. albicans*-treated HCT116 cells, and comparison of gap sizes (mean ± s.e.m.; n = 3), One-way ANOVA followed by Tukey’s multiple comparison test was used for comparisons. Groups labeled with any shared letters (e.g., “a” and “ab”, or “b” and “b”) indicate no significant differences were found between groups, while those without shared letters (e.g., “a” and “b”) indicate significant differences between groups. **(g)** Protein-level expressions of epithelial-mesenchymal transition (EMT) marker genes in HCT116 cells. **(h)** Immunofluorescence images showing the expression of EMT marker genes in HCT116 cells. White arrowhead represented fungal hypha. Scale bars represented 10 μm.

### C. albicans stabilizes HIF-1α and activates hypoxia response in CRC cells

To investigate the molecular mechanisms underlying *C. albicans*-induced cell migration, we performed whole-transcriptome sequencing (RNA-seq) on HCT116 cells co-cultured with *C. albicans.* As a result, we obtained significant up-regulation of 173 genes, which were potentially induced by *C. albicans* infection (Figure 2a, Suppl. Table S1). Of note, various pro-tumor cytokines, including TNF, CXCL8, CCL22, and IL6 family gene, as well as their corresponding receptors, were found to be up-regulated (Figure 2a,k, Suppl. Tables S2,S3), suggesting that *C. albicans* may actively contribute to the proinflammatory tumor microenvironment, thereby facilitating cancer progression^30^. Functional annotations of the up-regulated genes revealed enrichment in multiple metabolic pathways associated with tumor development, including hypoxia response, angiogenesis, epidermal growth factor signaling, and glucose metabolism (Suppl. Figure S2a and Table S4). Gene Set Enrichment Analysis (GSEA) further highlighted the up-regulation of the HIF-1α signaling pathway (Figure 2b, Suppl. Table S5). The up-regulation of key genes involved in HIF-1α signaling pathway, including HK2, LDHA, PDK1, and PFKFB1, were confirmed by qRT-PCR (Figure 2a,c, Suppl. Figure S2b). Moreover, treatment of HCT116 cells with L-mimosine, a compound that stabilizes HIF-1α, also significantly upregulated the expression of VEGFA, HK2, LDHA, PDK1, and PFKFB4 genes (Suppl. Figure S2c, S3b). To reinforce the findings, we cultured HCT116 cells under hypoxia condition and performed RNA-seq, revealing that 83.8% (145 genes) of the genes up-regulated by *C. albicans* were also upregulated under hypoxia (Suppl. Figure S2d).

**Figure 2.**
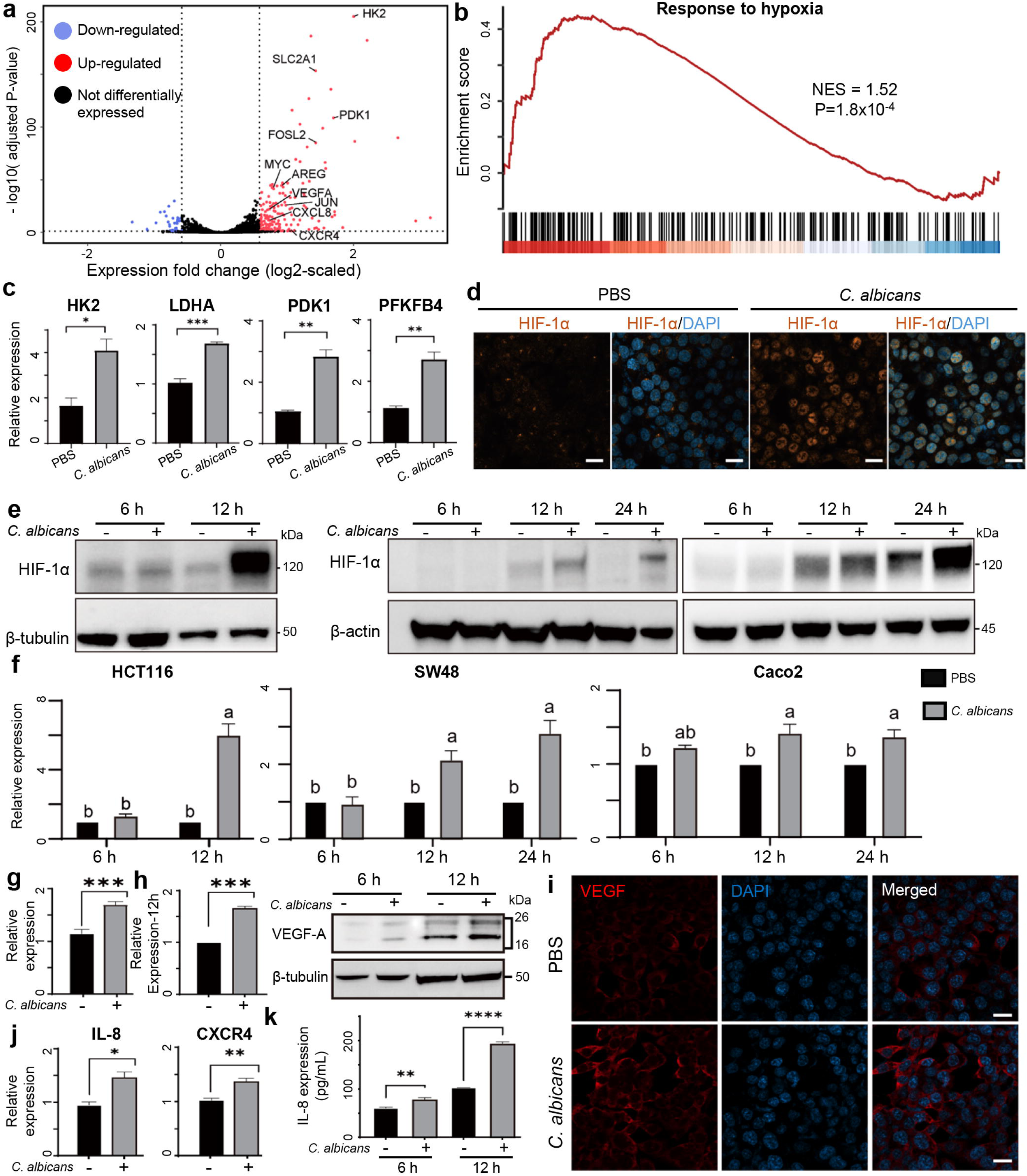
*C. albicans* induces hypoxia response pathway of colorectal cancer cells. **(a)** Differentially expressed genes in HCT116 cells after *C. albicans* infection for 12 hours versus PBS control, and **(b)** Gene Set Enrichment Analysis (GSEA) analysis showing up-regulation of “response to hypoxia” signaling pathway after *C. albicans* infection. **(c)** RNA-level expression of glycolysis genes in HCT116 cells cultured with *C. albicans.* **(d)** Immunofluorescence images showing HIF-1α expression after *C. albicans* infection of HCT116 cells. **(e,f)** Protein-level expression of HIF-1α in HCT116, SW48, and Caco2 cells after *C. albicans* infections. One-way ANOVA followed by Tukey’s multiple comparison test was used for comparisons. **(g)** RNA-level, **(h)** Protein-level expression, and **(i)** immunofluorescence of VEGFA gene in HCT116 cells after *C. albicans* infection for 12 hours. **(j)** RNA-level expression of IL-8 and CXCR4 genes HCT116 cells after *C. albicans* infection. **(k)** Concentration of IL-8 protein in cell culture supernatant. In (c, g, h, j, k), expressions were shown as mean ± s.e.m. (n=3), and t-tests were used to compare the expression levels between *C. albicans* infection and PBS control. *p < 0.05, **p< 0.01, ***p< 0.001. Scale bars indicated 10 μm.

Although the transcription level was unaffected, we observed the accumulation of HIF-1α in the nucleus and cytoplasm as early as 3 hours post-infection with immunofluorescence staining (Figure 2d, Suppl. Figure S3a-c). Western blots analysis further demonstrated that after *C. albicans* infection, HIF-1α increased in HCT116 and another two colorectal cell lines, SW48 and Caco2, while the timing of upregulation varies among these three cell lines (Figure 2e,f). PHD-mediated ubiquitination is the primary mechanism regulating HIF-1α protein stabilization^31^, we observed that although the total level of HIF-1α increased, the ratio of its ubiquitinated form to total HIF-1α decreased. This indicates that C. albicans activates hypoxia responses by stabilizing HIF-1α protein in cancer cells (Suppl. Figure S3d,e). Angiogenesis, a key process in cancer metastasis regulated by vascular endothelial growth factor (VEGF) ^32^, is also influenced by HIF-1α. Consistently, qRT-PCR analysis indicated increased VEGFA expression following *C. albicans* treatment (Figure 2g). Western blot and immunofluorescence analyses further confirmed enhanced production of VEGFA protein in tumor cells co-cultured with *C. albicans* (Figure 2h,i, Suppl. Figure S3c). In addition to VEGFA, the expressions of chemokine C-X-C motif ligand 8 (CXCL8) and C-X-C motif chemokine receptor 4 (CXCR4) were both upregulated in tumor cells following exposure to *C. albicans* (Figure 2j,k). These findings suggested that *C. albicans* infection may modulate the tumor microenvironment by promoting angiogenesis. To evaluate the cell type specificity of this response, we further analyzed additional epithelial cell models, as shown in Figure 8. Collectively, these findings suggested that *C. albicans* infection induces the hypoxia response pathway in colorectal cancer cells, thereby facilitated angiogenesis and metabolism adaptation for cancer progression.

### C. albicans activated c-Myc to promote HIF-1α pathway

Besides the genes shown in Figure 2c, the transcriptome data also uncovered a significant up-regulation of AP-1 (Activator protein 1) family and c-Myc genes in response to *C. albicans* infection, which we validated using qRT-PCR (Figure 3a). Considering the important roles of AP-1 in cell differentiation, migration, and proliferation^33^, the expressions of c-Jun and c-Myc were further substantiated by Western blots. For c-Jun, expression levels increased at 12 hours post-infection in HCT116 cells and at 24 hours in SW48 cells following *C. albicans* infection (Figure 3b). Similarly, *C. albicans* infection significantly elevated the expression of c-Myc in both HCT116 and SW48 cancer cell lines at 12 hours post-infection (Figure 3c). Notably, a high multiplicity of infection (MOI) accelerated the cellular response, observable as early as 6 hours post-infection (Suppl. Figure S4a).

**Figure 3.**
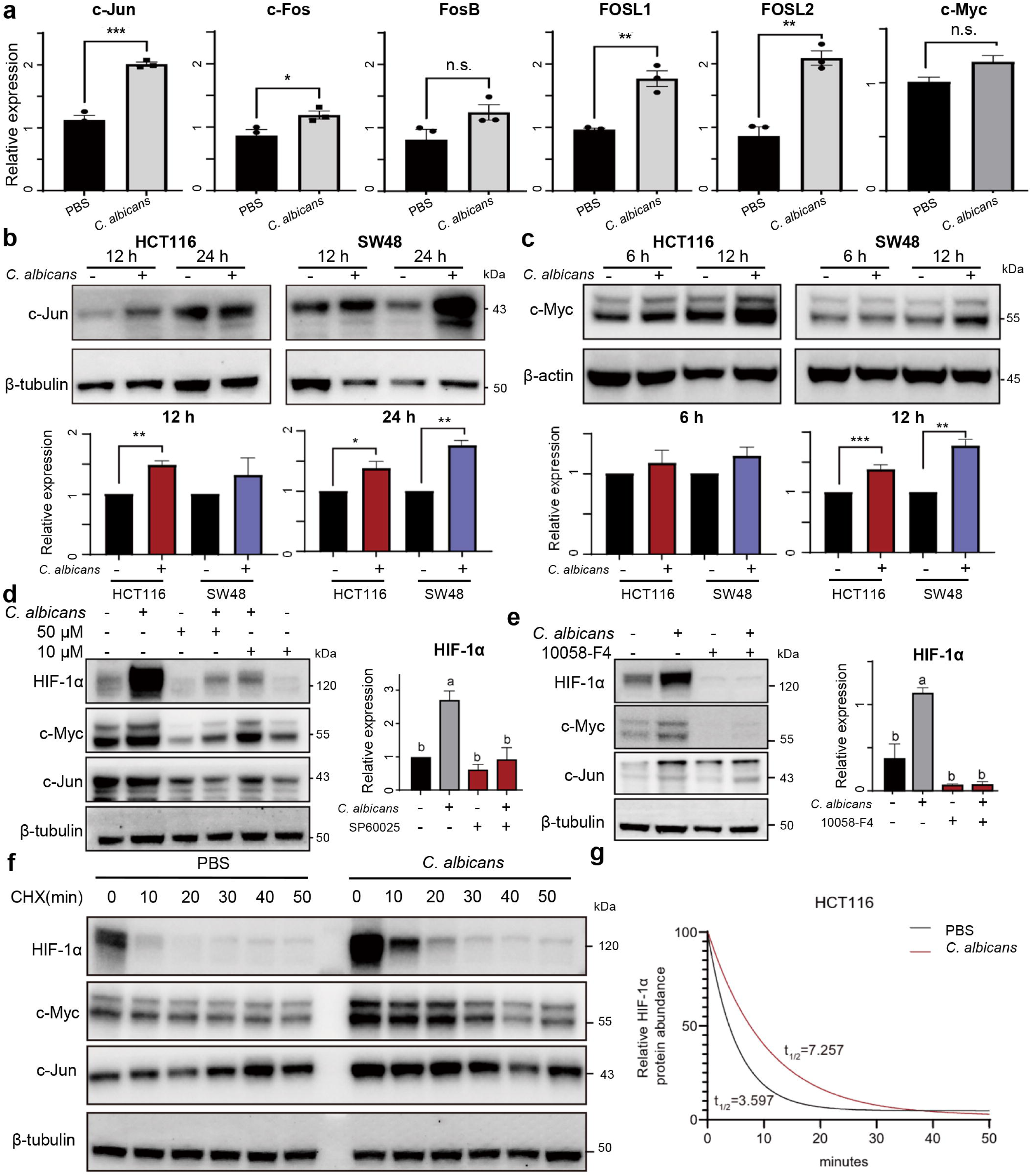
*C. albicans* up-regulate c-Myc and c-Jun to promote HIF-1α pathway. **(a)** RNA-level expression of AP-1 and c-Myc in HCT116 cells cultured with *C. albicans* for 12 hours. **(b)** Protein-level expressions of c-Jun in HCT116 and SW48 cells cultured with *C. albicans* for 12 and 24 hours. **(c)** Protein-level expressions of c-Myc in HCT116 and SW48 cells cultured with *C. albicans* for 12 and 24 hours. **(d)** Protein-level expressions of HIF-1α, c-Myc, and c-Jun in HCT116 cells after *C. albicans* infection and treated with c-Jun inhibitor SP600125 (with two dosage levels) for 12 hours. **(e)** Protein-level expressions of HIF-1α, c-Myc, and c-Jun in HCT116 cells after *C. albicans* infection and treated with c-Myc inhibitor 10058-F4 for 12 hours. **(f)** Protein-level expressions of HIF-1α, c-Myc, and c-Jun in HCT116 cells after *C. albicans* infection and treated cycloheximide. **(g)** The fitted curve of HIF-1α relative abundance after cycloheximide treatment. In (a, b), t-tests were used to compare the expression levels after *C. albicans* infection versus PBS control; in (d, e), one-way ANOVA followed by Tukey’s multiple comparison tests were used to compare various groups. n.s.: p>=0.05, *p < 0.05, **p< 0.01, ***p< 0.001.

To determine the regulatory hierarchy in the up-regulations of c-Jun, c-Myc, and HIF-1α, we constructed HIF-1α knock down cell line and found that *C. albicans* still robustly upregulated c-Jun and c-Myc levels (Suppl. Figure S4b). L-mimosine upregulated c-Jun but not c-Myc, suggesting a bidirectional regulatory relationship between c-Jun and HIF-1α (Suppl. Figure S4c). Treatment of HCT116 cells with the c-Jun inhibitor SP600125 or knock down of c-Jun expression inhibited the cellular accumulation of HIF-1α after *C. albicans* infection (Figure 3d, Suppl. Figure S4d,e); similarly, treatment with the c-Myc inhibitor 10058-F4 or KJ-Pyr9 inhibited the cellular accumulation of HIF-1α after *C. albicans* infection (Figure 3e, Suppl. Figure S4f). The results suggested that c-Jun and c-Myc act upstream in the regulatory cascade leading to HIF-1α up-regulation. Moreover, only inhibition of c-Myc with 10058-F4 or KJ-Pyr9 suppressed HIF-1α downstream gene expression following *C. albicans* infection (Suppl. Figure S4g). While c-Myc was reported to post-transcriptionally induce HIF-1α protein and target gene expression^34^, our time-course experiments with cycloheximide treatment further revealed that *C. albicans* infection may significantly stabilizes HIF-1α protein through c-Myc (Figure 3f-g, 4e-f). These finding indicate that c-Myc is the main regulator to mediate the cell hypoxia response to *C. albicans*.

### EGFR and TLR are involved in C. albicans-mediated cell response

Next, we aimed to identify the cell membrane receptor responsible for *C. albicans*-mediated response. Specifically, genes related to the regulation of EGF-activated receptor activity were enriched following *C. albicans* infection (Suppl. Figure S2a). Indeed, both RNA-seq and qRT-PCR confirmed that *C. albicans* induced the expression of amphiregulin (AREG), a member of the epidermal growth factor (EGF) family and a known ligand of EGFR (Figure 4a). To explore the role of EGFR in the response to *C. albicans*, we treated HCT116 cells with the EGFR kinase inhibitor AG-1478. As a result, AG-1478 significantly reduced the activation of c-Myc and half-lives of HIF-1α after *C. albicans* infection, indicating inhibition of the response triggered by *C. albicans* (Figure 4b-f); Given the complex regulatory network between HIF-1α, c-Jun, c-Myc and the mitogen-activated protein kinase (MAPK) pathway^34–40^, we investigated the phosphorylation statuses of Erk and P38 (gene symbol: MAPK14), two important compartments of the EGFR-MAPK pathway. Western blots analysis revealed that *C. albicans* infection significantly increased the phosphorylation of Erk (Figure 4g), and treatment with the ERK inhibitor SCH772984 effectively suppressed the phosphorylation of Erk, consequently inhibiting the cellular responses triggered by *C. albicans*. On the other hand, phosphorylation of P38 remained unaffected after *C. albicans* infection, and treatment with the P38 inhibitor adezmapimod did not produce any significant inhibitory effect (Figure 4h), indicating a selective role of the ERK pathway in the cellular response to *C. albicans* infection.

**Figure 4.**
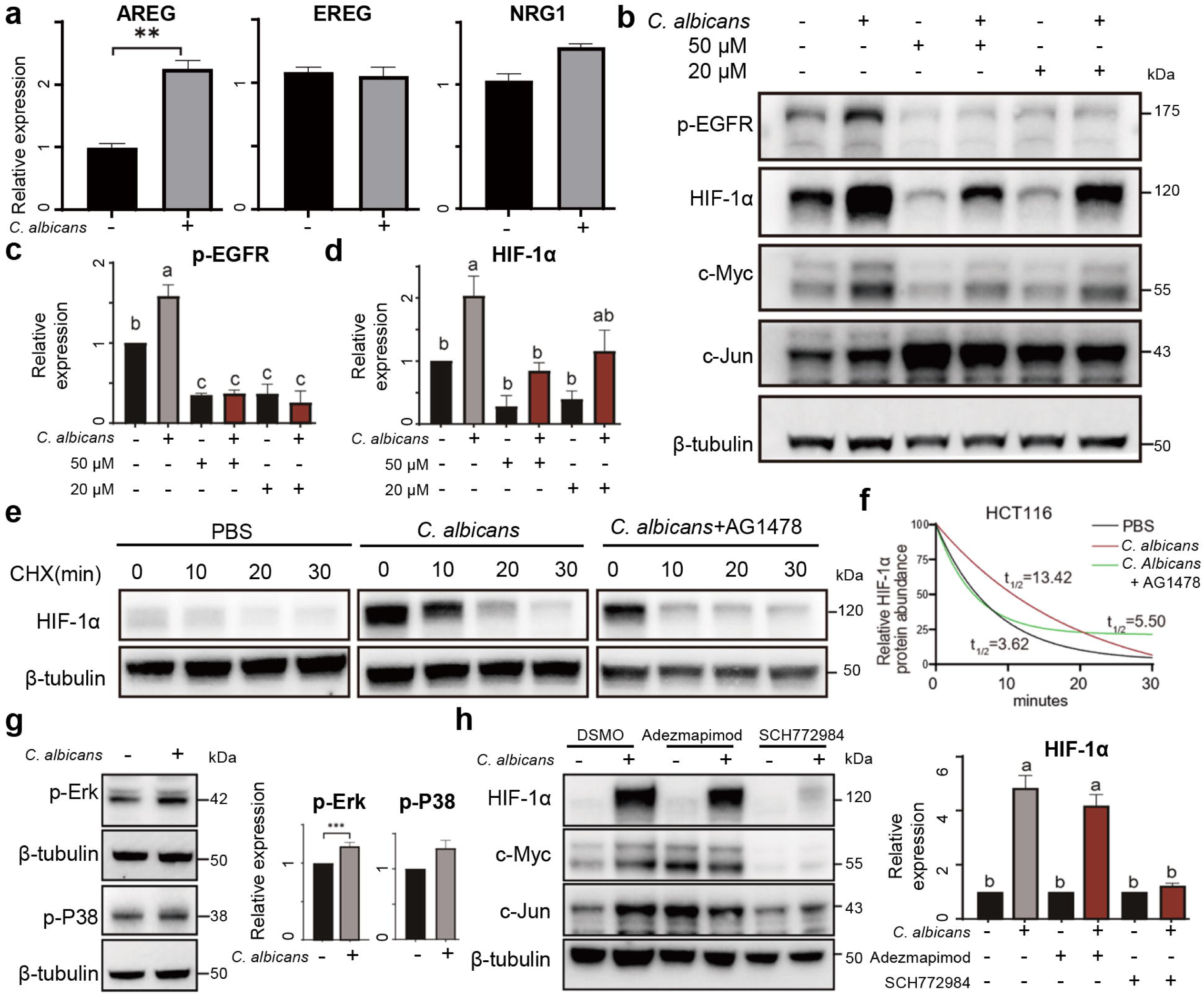
EGFR-ERK is involved in *C. albicans*-mediated cancer cell response. **(a)** RNA-level expressions of EGFR ligands in HCT116 cells cultured with *C. albicans* for 12 hours. **(b-d)** Protein-level expressions of p-EGFR, HIF-1α, c-Myc, and c-Jun in HCT116 cells treated with EGFR inhibitor AG-1478 (with two dosage levels) for 12 hours. **(e)** Protein-level expressions of HIF-1α in HCT116 cells after *C. albicans* infection and treated with cycloheximide. **(f)** The fitted curve of HIF-1α relative abundance after cycloheximide treatment. **(g)** Protein-level expressions of p-Erk and p-P38 in HCT116 cells after *C. albicans* infection. **(h)** Protein-level expressions of HIF-1α, c-Myc, and c-Jun in HCT116 cells treated with P38 inhibitor adezmapimod or Erk inhibitor SCH772984 for 12 hours. In (a,e), expressions between *C. albicans* infection and PBS control were compared using t-tests; in (c,d,f), one-way ANOVA followed by Tukey’s multiple comparison tests were used to compare various groups. **p< 0.01, ***p< 0.001.

Notably, despite treatment with AG1478, the levels of HIF-1α and c-MYC in *C. albicans*-infected cells remained significantly elevated compared to the uninfected control group (Figure 4b). We hypothesized that this residual effect might be due to the continued activation of cell surface Toll-like receptors (TLRs) or C-type lectin receptors (CLRs). Despite the low expression of TLR and CLR in HCT116 cells (Suppl. Figure S5a, S6a), we successfully generated stable knockdown cell lines for key receptors and pathways associated with fungal recognition, including TLR2, TLR6, TLR9, MYD88, Dectin-1, and SYK (Suppl. Figure S5b). Western blot results showed that neither the suppression of genes nor treatment with the TLR2 inhibitor MMG11 or the Dectin1 inhibitor laminarin impaired the cell responses following *C. albicans* infection (Figure 5a, Suppl. Figure S5c-e, S6b-d). However, when cells were treated with a combination of receptor inhibitors targeting both EGFR and TLR2, the activations of HIF-1α and c-MYC induced by *C. albicans* were completely suppressed (Figure 5b). Furthermore, we investigated the status of NF-κB, a key pathway responsive to TLR activation, after *C. albicans* infection^41–43^. Both immunofluorescence staining and western blot analyses revealed an increase in the phosphorylation level of p65 (gene symbol: RELA), a principal subunit of the NF-κB complex, following *C. albicans* infection. This led to nuclear translocation of p65, which modulated the expression of several downstream genes, including c-Jun and HIF-1α (Figure 5c-g). Additionally, combined EGFR and TLR2 inhibitors completely reversed the enhanced cell migration induced by *C. albicans* (Figure 5h,i). Collectively, the results suggested that *C. albicans* modulated the intracellular stability of HIF-1α by activating EGFR-ERK and TLR-NF-κB pathways, which further up-regulated a cohort of cellular transcription factors, including AP-1 components and c-Myc.

**Figure 5.**
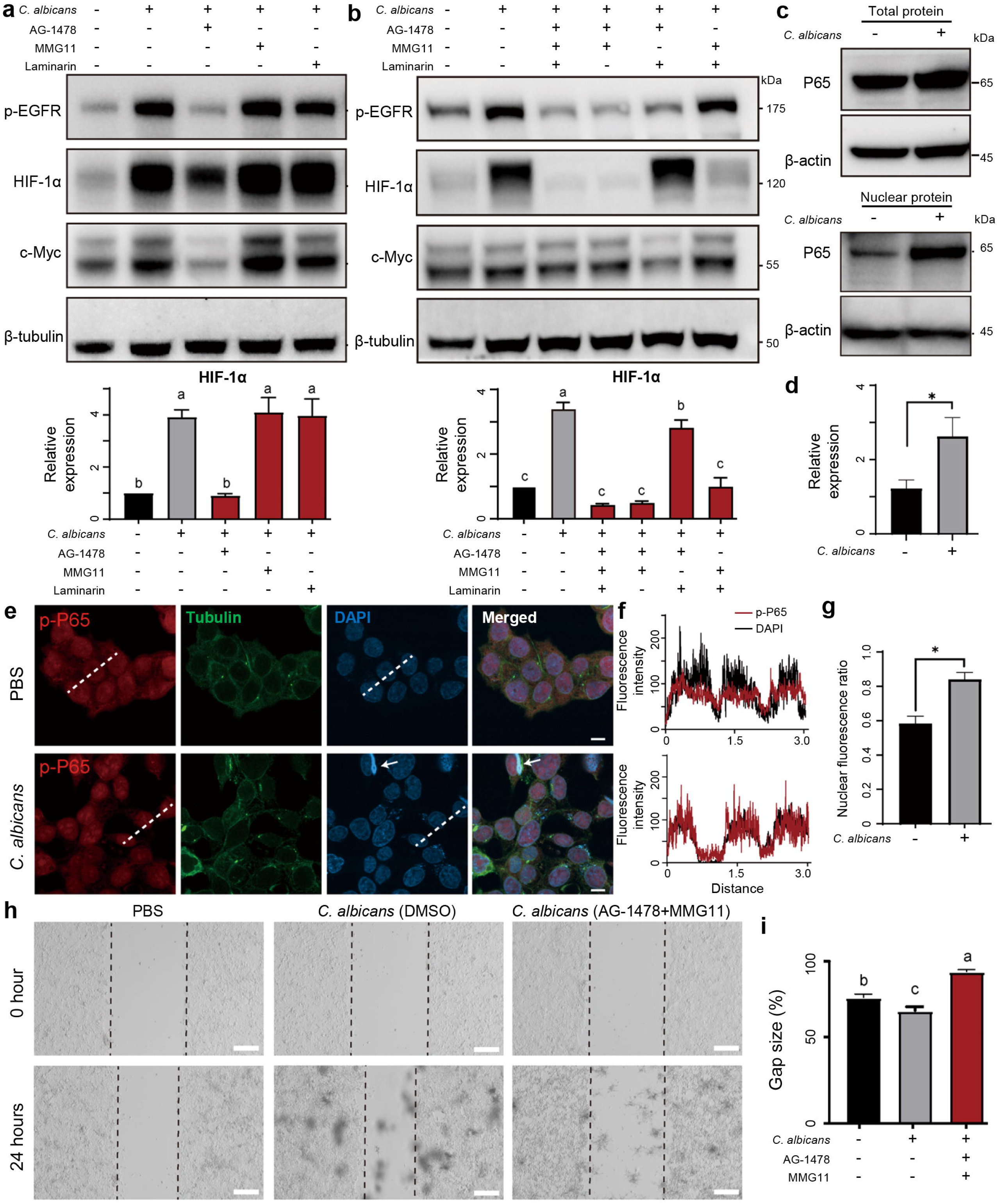
*C. albicans* modulated the stability of HIF-1α by activating EGFR-ERK and TLR-NF-κB pathways. **(a,b)** Protein-level expression of p-EGFR, HIF-1α, and c-Myc in HCT116 cells after *C. albicans* infections with AG-1478/MMG11/Laminarin treatments. **(c)** Expression of p-P65 in HCT116 cells (whole cell or nucleus), and **(d)** comparison of p-P65 protein level in HCT116 nuclei cultured with *C. albicans* for 12h. **(e)** Immunofluorescence images showing p-P65 localization, **(f)** relative signal intensities of the respective emission fluorescence along the lines drawn in (e), and **(g)** comparison of nuclear fluorescence ratios in HCT116 cells after *C. albicans* infection. Scale bars represented 10 μm and were applied across all images in this panel. **(h)** Wound-healing assay measuring migration capability of HCT116 cells after *C. albicans* infection with or without AG-1478/MMG11 treatment, and **(i)** comparison of gap sizes of different groups. In **(d, g)** t-tests were used to compare the two groups; in **(a,b,i),** one-way ANOVA followed by Tukey’s multiple comparison tests were used. *p < 0.05.

### Candidalysin is essential for eliciting colorectal cancer cell responses

To explore the virulence factor responsible for inducing cell responses, we co-cultured *C. albicans* with HCT116 cells and harvested *C. albicans* at various time points post-infection. qRT-PCR assays revealed that Als3, Hwp1, and Ece1 genes were significantly up-regulated in post-infection *C. albicans* (Figure 6a). Als3 and Hwp1 are known to be involved in the adhesion of *C. albicans* to epithelial cells, while Ece1 encodes the virulence factor candidalysin^21^. Interestingly, the expression level of Ece1 showed a >1,000-fold increase (Figure 6a-b). In fact, during the early stages of mucosal surface infection by *C. albicans*, candidalysin is released into the invasion pocket created by the penetrating hyphae^21^, where it activates danger-response and damage-protection pathways in host cells, including the activation of the epidermal growth factor receptor (EGFR) in epithelial cells and the NLRP3 inflammasome in macrophages^44,45^. Hence, these results suggest that candidalysin might play a critical role interacting with HCT116 cells.

**Figure 6.**
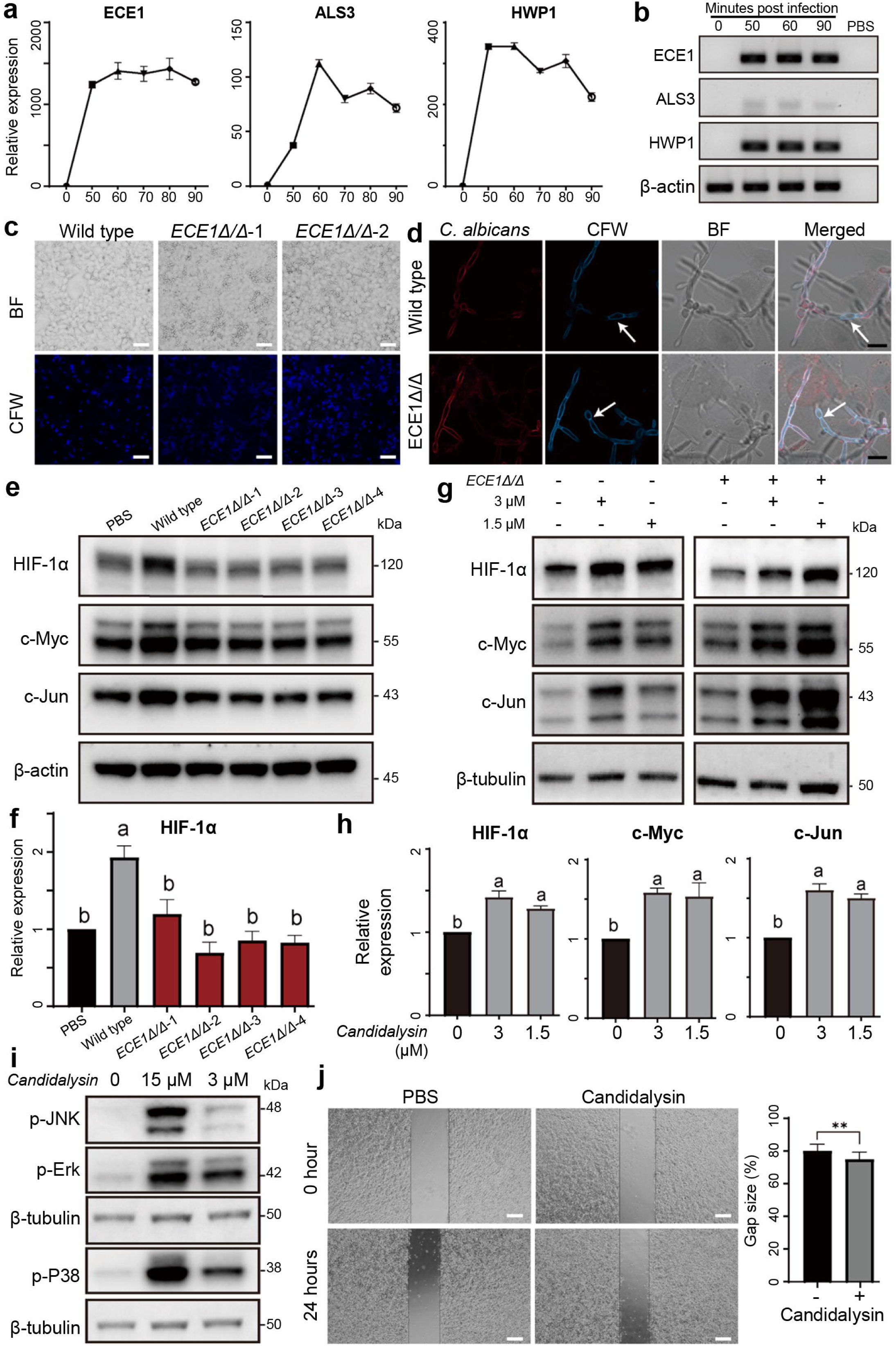
Candidalysin is critical for eliciting colorectal cancer cells response. **(a, b)** Expressions of CaECE1, CaALS3 and CaHWP1 cultured with HCT116 cells at 0 to 90 minutes post infection measured by (a) qRT-PCR and (b) RT-PCR. **(c)** Fluorescence staining of *C. albicans* wild-type and ECE1Δ/Δ hyphae adherent to HCT116 cells. Scale bars were 50 μm. **(d)** Fluorescence staining of *C. albicans* wild-type and ECE1Δ/Δ hyphae invading through HCT116 cells. White arrowheads indicate hyphal invading the cells. Scale bars were 10 μm. **(e, f)** Protein-level expressions of HIF-1α, c-Jun, and c-Myc in HCT116 cells co-cultured with wildtype or ECE1Δ/Δ strain *C. albicans* for 12 hours. **(g, h)** Protein-level expressions of HIF-1α, c-Jun, and c-Myc in HCT116 cells co-cultured with ECE1Δ/Δ strain *C. albicans* and treated with or without candidalysin. **(i)** Activation of MAPK pathway in HCT116 cells treated with candidalysin. **(j)** Wound-healing assay for migration capability of HCT116 cells treated with candidalysin (3 μM). In **(f, h)**, ANOVA followed by Tukey’s multiple comparison test was used to compare different groups. In (j), t-tests were used to compare the two groups; **p< 0.01.

To investigate the function of candidalysin during *C. albicans*-colon cell interaction, we generated an Ece1 knockout strain (termed as “Ece1Δ/Δ” hereafter) using the CRISPR/Cas9 system in *C. albicans* (Suppl. Figure S7a-b). This strain, which cannot synthesize candidalysin, was assessed for its ability to penetrate HCT116 cells using a differential fluorescence assay^21,46^. The Ece1Δ/Δ strain maintained the capacity to adhere to and enter HCT116 cells, similarly to the wild-type strain (Figure 6c-d). However, it failed to activate HIF-1α and c-Myc in the tumor cells (Figure 6e,f). In contrast, when HCT116 cells were treated with synthetic candidalysin, we observed a significant upregulation of c-Jun, c-Myc, and HIF-1α expression levels at 3μM for both uninfected HCT116 cells and those infected by Ece1Δ/Δ strain *C. albicans* (Figure 6g,h). Moreover, treated with candidalysin activated MAPK pathway and showed promoted cell migration (Figure 6i, j). Together, these results indicate that *C. albicans* regulates tumor cells through the secretion of candidalysin, which activates MAPK and hypoxia pathways to promote tumor cell migration.

### C. albicans induce hypoxia responses in colon organoids and clinical specimens

To investigate whether *C. albicans* has the same effect on tumor samples, we collected three paired tumors and non-malignant colon tissues originating from CRC patients, which were isolated and cultured into organoids. After co-culturing *C. albicans* with the organoids for 12 hours, we also detected significant aggregation of HIF-1α in the cells, as evidenced by both Western blot and immunofluorescence staining, in both non-malignant intestinal organoids and tumor organoids (Figure 7a-c, Suppl. Figure S8a). Treatment of the organoids from colorectal cancer patients with c-Jun inhibitor SP600125 and c-Myc inhibitor 10058-F4 significantly suppressed the expression of HIF-1α (Figure 7d, Suppl. Figure S8b). Moreover, we observed that *C. albicans* hyphae penetrated the epithelium of colon organoids, reaching the lumen (Figure 7e). For the specimens analyzed in Figure 1a, we found higher levels of fungi near the colon mucosa in stage III samples compared to stage II sample, along with a notable aggregation of HIF-1α (Figure 7f, Suppl. Figure S9). Collectively, our findings suggest that *C. albicans* may contribute to the development of CRC by inducing HIF-1α aggregation in adjacent cells, thereby triggering hypoxia-related responses that create a microenvironment conducive to tumor progression.

**Figure 7.**
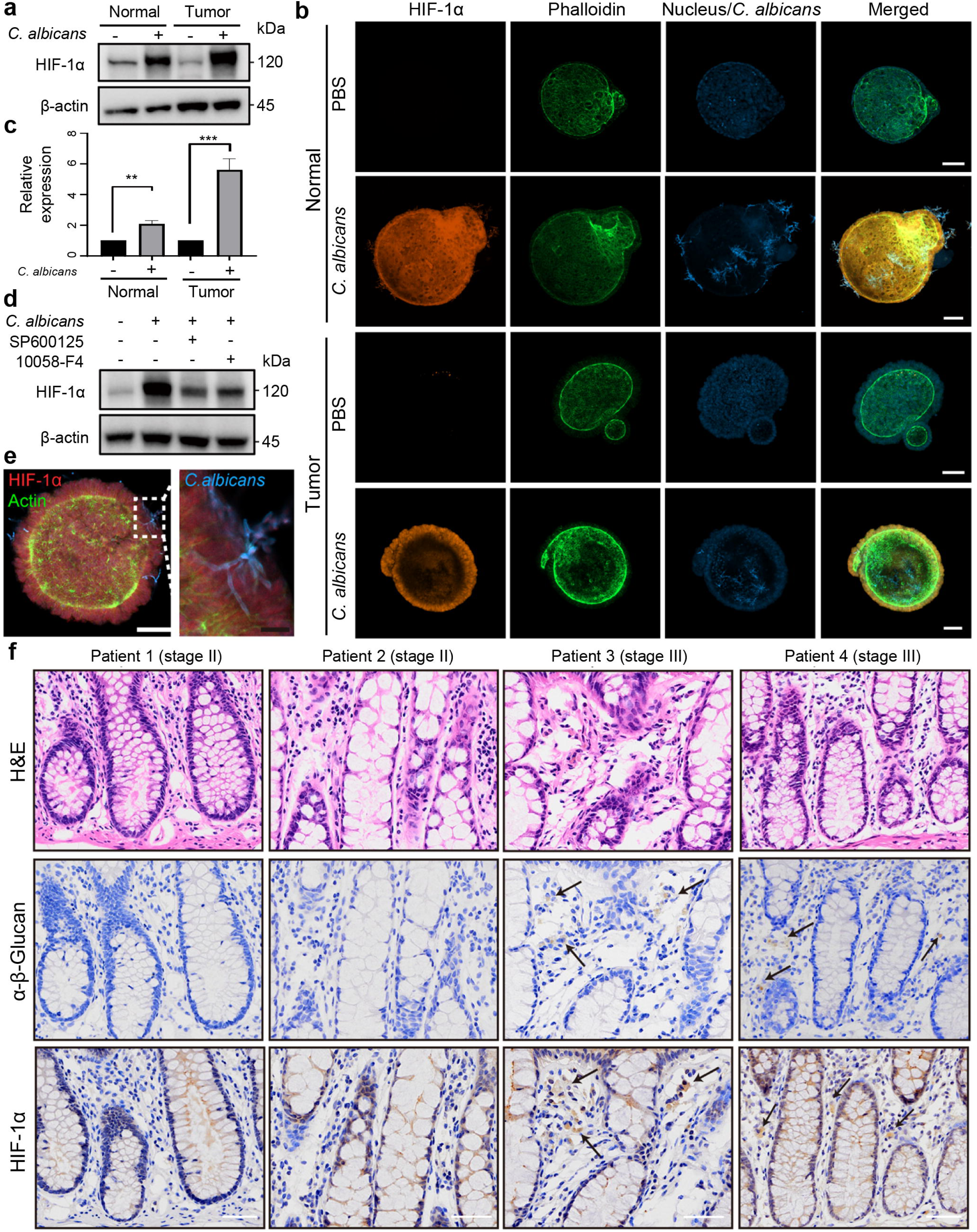
*Candida albicans* induces hypoxia response pathway of colon organoids and CRC samples. **(a,b)** Protein-level expression and immunofluorescence of HIF-1α in normal and CRC organoids after *C. albicans* infection. **(c)** Quantification of HIF-1α protein levels in (a). **(d)** HIF-1α expression after *C. albicans* infection and treated with c-Jun inhibitor SP600125 or c-Myc inhibitor 10058-F4 in CRC organoids. **(e)** Immunofluorescence images of CRC organoids after *C. albicans* infection. White and black scale bars represented 50 and 10 μm, respectively. **(f)** Representative histological images of tumor tissues stained by hematoxylin and eosin (H&E), antibodies against β-glucan and HIF-1α. Scale bars indicated 50 μm. In (c), t-tests were used to compare normal or tumor organoids after *C. albicans* infections versus PBS controls. **p< 0.01, ***p< 0.001.

### Interactions between Candida and tumor cells exhibit both species and cellular specificity

Lastly, we explored the cellular responses of CRC cells to other fungus besides *C. albicans.* To do this, we conducted co-culture experiments using HCT116 cells and two other fungal species*, Saccharomyces cerevisiae* and *Candida tropicalis*, to assess their adhesive capacities and effects on cancer cells. As shown in Figure 8a, *C. tropicalis* and *S. cerevisiae* exhibited minimal adhesion to cancer cells. Western blots were then performed to examine the subsequent effects of infection with these fungi, as well as a mixture of fungal cell wall polysaccharides and heat-killed Candida species. As a result, *C. albicans* and *C. tropicalis* exhibited the most potent ability to activate the hypoxia pathway. *C. albicans* induced an increase in c-Myc expression, with a trend toward higher levels compared to other fungal specie*s*. (Figure 8b). We hypothesized that the variations in cellular responses might be linked to the diversities of the Ece1 gene in the fungal genomes. Indeed, phylogenetic analysis revealed that evolutionarily, the Ece1 genes in various *Candida* species were highly conserved, whereas they differed significantly from other fungus. (Suppl. Figure S10). Of note, *C. tropicalis* did not express the ECE1 homolog after interaction with HCT116 cells for 1 hour (Suppl. Figure S11).

**Figure 8.**
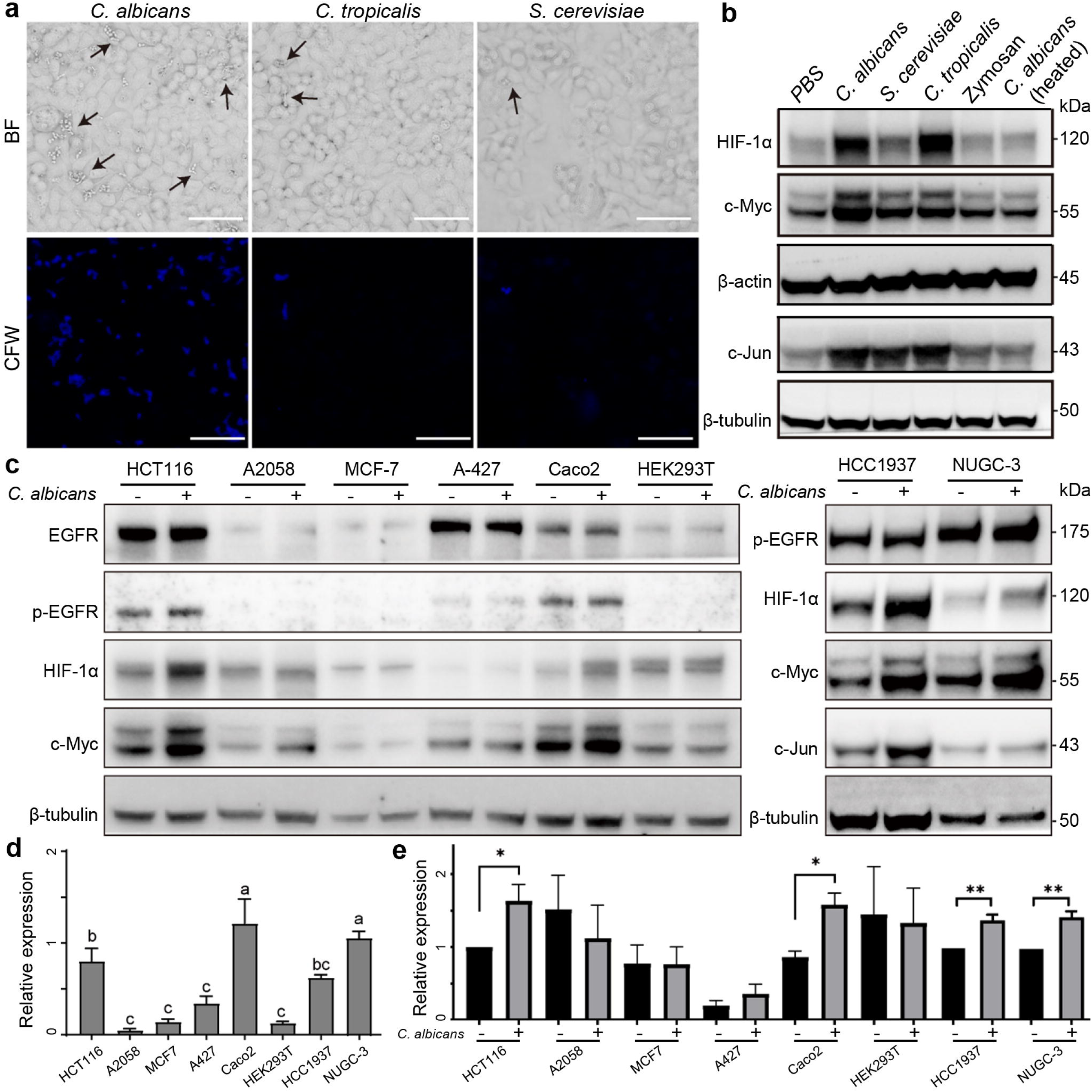
The interactions between Candida albicans and tumor cells exhibit both species and cellular specificity. **(a)** Fluorescence staining of *C. albicans*, *C. tropicalis,* and *S. cerevisiae* adhesion to HCT116 cells. Black arrowheads indicated yeast or hyphal adherent to HCC116 cells. Scale bars were 50 μm. **(b)** Protein-level expressions of HIF-1α, c-Myc, and c-Jun in HCT116 cells after infections of various types of microbiota. **(c)** Protein-level expressions of EGFR, p-EGFR, HIF-1α, and c-Myc in various cancer cells after *C. albicans* infections. **(d)** Quantifications of p-EGFR and **(e)** HIF-1α expressions in **(c)**. In (d), One-way ANOVA followed by Tukey’s multiple comparison test were used to compare across cells; in (e), t-tests were used to compare expressions between *C. albicans* infection and PBS control of the same cell type. *p < 0.05, **p< 0.01.

To further assess the cell type specificity of this response, we investigated the response to *C. albicans* by co-culturing it with a variety of cancer cell lines. Intriguingly, we found that *C. albicans* not only activated the hypoxia pathway in CRC cells but also significantly induced similar responses in breast cancer cell line HCC1937 and the gastric carcinoma cell line NUGC-3. In contrast, *C. albicans* failed to elicit responses in melanoma cell line A2058, the breast ductal carcinoma cell line MCF-7, and the human embryonic kidney cell line 293T (HEK293T) (Figure 8c). To explore the underlying mechanism responsible for these differences, we analyzed EGFR expression. As shown in Figure 8d, e, Suppl. Figure. S12, the results indicated a correlation between the expression and phosphorylation levels of EGFR and the phenotypic response to *C. albicans* infection. This was consistent with our earlier finding that *C. albicans* facilitates tumor cell migration through an EGFR-mediated pathway (Figure 4). Together, these findings suggest that candidalysin triggered EGFR/TLR2 - ERK/NF-κB - HIF-1α signaling pathway to induce hypoxia response, which promotes the progression of CRC (Figure 9), and these observations support a context-dependent response to *C. albicans* across epithelial cell types.

**Figure 9.**
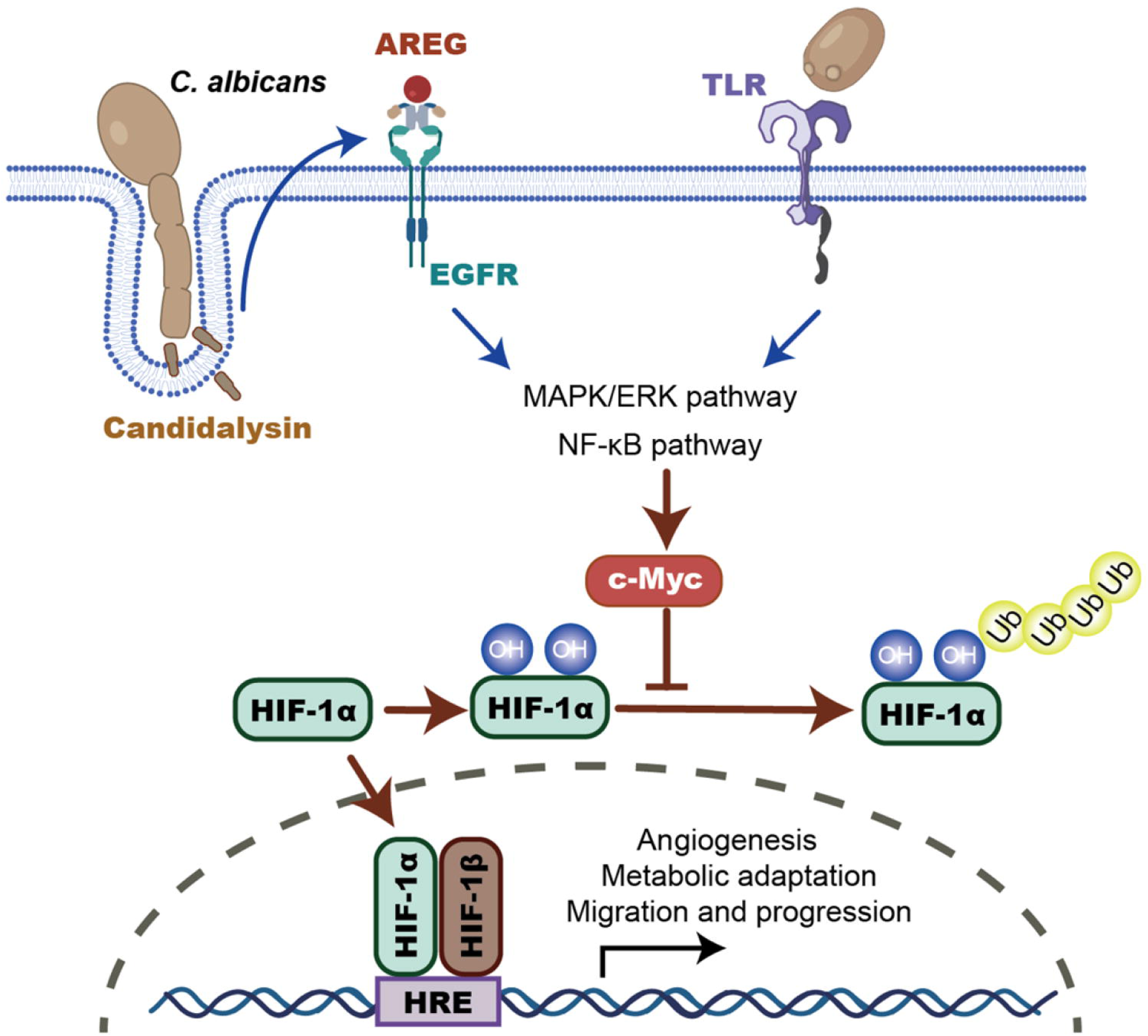
Schematic diagram of the *C. albicans* infection during colorectal cancer progression. This diagram depicts the proposed signaling pathway activated by *C. albicans* through the secretion of its peptide toxin, candidalysin. Upon interaction with colorectal cancer (CRC) cells, *C. albicans* engages epidermal growth factor receptor (EGFR) and toll-like receptor 2 (TLR2), triggering downstream signaling cascades. These interactions result in the activation of the mitogen-activated protein kinase (MAPK) pathway and nuclear factor kappa B (NF-κB), both of which are key mediators of cellular stress and inflammatory responses. The coordinated activation of these pathways leads to stabilization and accumulation of hypoxia-inducible factor 1-alpha (HIF-1α) through c-Myc, thereby inducing a hypoxic transcriptional program even under normoxic conditions. This hypoxia-like response contributes to the pro-tumorigenic microenvironment and may promote colorectal cancer progression.

## Discussion

Recent studies have sparked broad interest in the role of gut fungi, including *C. albicans*, in the development of various types of cancers^24,26,47^. However, whether and how fungal components directly modulate epithelial oncogenic signaling remains largely unexplored. We identify *C. albicans* induce hypoxia response in colorectal cancer (CRC) cells, mediated by the activation of the EGFR/TLR2–ERK/NF-κB signaling axis and stabilization of HIF-1α. This process is critically dependent on the fungal virulence factor candidalysin. While prior research has highlighted fungal modulation of tumor-associated macrophages^26^, our study extends this concept by identifying direct fungal–epithelial crosstalk driving oncogenic signaling in CRC. Hence, by demonstrating that *C. albicans* activates epithelial hypoxia signaling in a species- and cell-specific manner, our study reveals a previously unrecognized mechanism through which gut fungi may contribute to CRC progression.

Hypoxia is a hallmark of cancer, promoting angiogenesis and metastasis and resistance to therapy^48–50^. Under hypoxic conditions, metabolic shift in malignant cells enhance cell proliferation and invasion ability^51^, as well as induce EMT and modulate the tumor microenvironment^52,53^ to promote metastasis^54^. Interestingly, previous studies have shown that tumor-associated macrophages with *C. albicans* infection showed an increased glycolysis level through the HIF-1α-dependent pathway^26^. Similarly, it has been reported that *C. albicans* can upregulate the mRNA expression of HIF1A and CAMP in HT-29 colonocytes^27^. In our study, however, we found that the *C. albicans*-induced hypoxia response in colorectal cancer (CRC) cells is predominantly regulated at the protein level, indicating post-transcriptional or translational control mechanisms. Moreover, we demonstrated that this hypoxia response relies on c-Jun and c-Myc, two key cancer-associated transcription factors that regulate cell progression, apoptosis, and cellular transformation^55,56^. This finding aligns with recent reports showing that c-Myc exerts considerable control over the levels and activity of HIF-1α^36,57^. Besides hypoxia, *C. albicans* also induced VEGFA expression (Figure 2), suggesting that the fungus may contribute to the vascularization of tumors, thereby facilitating nutrient supply and waste removal to boost tumor growth. Furthermore, activation of the NF-κB pathway by *C. albicans* suggested a potential role in modulating the inflammatory response worthwhile for future investigations.

In addition, we demonstrated that candidalysin is the key molecular that triggers CRC cell response after *C. albicans* infection. Candidalysin is a peptide toxin secreted by the hyphal form of *C. albicans* during invasion, which helps *C. albicans* compete with bacterial species in the intestinal niche^58^, and it is known to drive epithelial damage, immune activation, and phagocyte attraction^21,44,45,59^. Recent studies have also shown that candidalysin can modulate multiple signaling pathways in various cancer types, e.g., activate the MAPK p38 mitogen-activated protein kinase through MAPK kinases and the kinase Src in oral cancer^59^. Here we showed that candidalysin acted as a critical virulence factor that induced phosphorylation of EGFR to activate MAPK/ERK and downstream HIF-1α pathway. Together, these findings suggested that candidalysin, and its downstream EGFR, could serve as therapeutic targets to suppress *C. albicans*-induced metastasis-promoting responses for pharmacological interventions. Notably, we also tested additional epithelial models, including normal colon-derived organoids and breast epithelial cancer cells, which did not exhibit HIF-1α activation upon *C. albicans* exposure, suggesting that this response is not a universal epithelial phenomenon. (Figure 8). This variability may be influenced by factors such as EGFR expression levels or other epithelial-specific signaling determinants. However, given the limited number of models tested, further studies are required to systematically evaluate species- and cell type-specific responses. Interestingly, *C. tropicalis* are able to induce HIF-1α accumulation, while it does not secret candidalysin (Figure 8b, Suppl. Figure S11)^60^. In fact, previous studies reported that *C. tropicalis* induce PKM2 phosphorylation which then upregulates HIF-1α pathway^61^, suggesting high diversity of fungus-tumor interplays that requires more explorations. Inhibition of TLR2 resulted in a stronger effect on downstream signaling than inhibition of Dectin-1, despite the established role of Dectin-1 in recognizing *C. albicans* β-glucans. One possible explanation is that the expression level of Dectin-1 is relatively low in these epithelial cancer cells, thereby limiting its functional contribution. In addition, differences in ligand accessibility, particularly in the context of fungal cell wall architecture, may influence receptor engagement. It is also possible that redundancy and cross-talk among pattern recognition receptor pathways compensate for Dectin-1 inhibition. These observations highlight the context-dependent nature of host–fungal interactions.

A key limitation of this study is the small number of clinical samples analyzed, which restricts the strength of conclusions regarding the association between *C. albicans* and CRC progression in patients. Therefore, future studies with larger and well-characterized patient cohorts will be required to validate these findings and determine their clinical relevance. Another limitation of this study is the lack of *in vivo* validation. While our findings are supported by multiple in vitro models and patient-derived organoids, *in vivo* studies will be necessary to fully establish the role of *C. albicans* in CRC progression within a physiological context. Notably, previous studies have reported that *C. albicans* colonization promotes tumor burden and inflammation in mouse models, supporting the biological relevance of our findings. Future work integrating *in vivo* models will be essential to validate and extend the mechanistic insights described here.

In conclusion, this study reveals a novel link between fungal infection and hypoxia activation in the tumor microenvironment. Given the central role of HIF-1α in metabolic adaptation, angiogenesis, and therapeutic resistance, our study provides a promising framework for understanding how microbial factors may influence CRC progression through metabolic reprogramming.

## Methods

### Ethics approval and cell culture

This study had been approved by the Ethics Committee of Shenzhen Bay Laboratory, Ethics Committee of Southern University of Science and Technology, and Ethics Committee of The Seventh Affiliated Hospital, Sun Yat-Sen University. Clinical colon tumor samples were collected from Southern University of Science and Technology Hospital and The Seventh Affiliated Hospital, Sun Yat-Sen University. The human CRC cell line HCT116 (RRID: CVCL_0291), gastric adenocarcinoma cell line NUGC-3 (RRID: CVCL_1612), breast cancer cell line HCC1937 (RRID: CVCL_0290) and non-small cell lung cancer A-427 (RRID: CVCL_1055) cultured in RPMI 1640 Medium (Thermo, #C11875500BT), supplemented with 10% Fetal Bovine Serum (FBS) (Sigma-Aldrich, #F0193), at 37°C in a humidified incubator with an atmosphere of 5% CO_2_. The human CRC cell lines SW48 (RRID: CVCL_1724) and Caco2 (RRID: CVCL_0025), breast cancer cell line MCF-7 (RRID: CVCL_0031), melanoma cell line A2058 (RRID: CVCL_1059), and embryonic kidney cells HEK293T (RRID: CVCL_0063) are grown in Dulbecco’s Modified Eagle Medium (DMEM) (Thermo, #C11995500BT) supplemented with 10% FBS, under the same conditions. For detected gene expression under hypoxia, HCT116 cells were cultured at 37□ in an anaerobic chamber with an atmosphere of 0% O_2_, 5% CO_2_ and 95% N_2_. Cells were routinely tested for mycoplasma contamination and were found to be negative. *C. albicans* (RRID: NCBITaxon_5476), *C. tropicalis* (RRID: NCBITaxon_5482), and *S. cerevisiae* (RRID: NCBITaxon_4932) were grown on YPD (Yeast Peptone Dextrose) agar plate (HKM, #21100). Prior to infection, a single colony was inoculated into 4 mL YPD liquid medium and incubated overnight at 30°C in a high-humidity environment. *Bacteroides fragilis* was cultured on columbia blood agar plate (HKM, #CP0160) and inoculated with a single colony into 5 mL BHI broth, supplemented with 25μg/mL hemin chloride and 10μg/mL vitamin K1(HKM, #SR0300). Cultures were maintained in an anaerobic chamber at 37□.

### Co-culture of cells and strains

For co-culture of cell line and strains, 1×10^6^ cells were grown to confluence on 6-well plate. Before co-culture, the medium was replaced with 2 mL RPMI 1640 without FBS and supplemented with the strains, then incubated for 6 or 12 hours at 37 °C/5% CO_2_. For co-culture of organoids and *C. albicans*, approximately 200 colon organoids (∼5×10^5^ cells) were harvest from a 12-well plate. Organoids were gently washed with 1 mL of DMEM medium without FBS, then the DMEM was aspirated and replaced with 1 mL of the organoid culture medium. The medium was then replaced with 1 mL serum-free medium supplemented with *C. albicans* and incubated for 12 hours at 37 °C/5% CO_2_. Unless otherwise specified, the MOI (multiplicity of infection) used in the infection experiments was set at a ratio of strain: cell = 1:25.

### Cell migration assay

Cell migration was assessed using a wound-healing assay. Briefly, HCT116 cells were seeded in 6-well culture plates to reach approximately 90% confluence. A wound was introduced by manually scraping. Following the initial wound creation, cells were washed with phosphate-buffered saline (PBS) to remove debris and 2 mL serum-free medium supplemented with the strains was added. Cells were then incubated at 37°C in a CO_2_ incubator for 24 hours. Images of the wounded area were captured at 0 and 24 hours, and the gap sizes were quantified by measuring the wound area using ImageJ software. The relative gap size was calculated by comparing the wound area at 24 hours to the initial wound size.

### RNA extraction and real-time PCR

Total RNA was isolated from cultured tumor cells using a commercial kit (Zymo, #R2052), followed by quantification and purity assessment with nanodrop. 1μg of total RNA were treated with DNase to eliminate DNA contamination and then converted to cDNA using a high-capacity cDNA synthesis kit (Vazyme, #R333-01). For quantitative Real-Time PCR (qPCR), gene-specific primers were used in a SYBR Green-based qPCR assay to quantify the expression levels of target genes (Vazyme, #Q712-02). The qPCR cycling conditions included an initial denaturation step, followed by 40 cycles of amplification. Relative gene expression was determined using the 2^-ΔΔCt^ method with β-actin served as endogenous control for normalization. The DNA primers were listed in Suppl. Table S6.

### Protein extraction and western blots

Tumor cells were lysed in a radioimmunoprecipitation assay (RIPA) buffer (Beyotime, #P0013) containing protease and phosphatase inhibitors (Selleck, #B14001, #B15001). The lysate was centrifuged, and the supernatant containing the total protein was collected. Protein concentration was determined using the detergent compatible bradford protein assay kit (Beyotime, #P0006C) with bovine serum albumin (BSA) used for standard curve construction. For western blot analysis, 10μg protein samples were separated by sodium dodecyl sulfate-polyacrylamide gel electrophoresis (SDS-PAGE) and then transferred to a polyvinylidene fluoride (PVDF) membrane. After blocking with 5% non-fat dry milk for one hour, the membrane was incubated with a primary antibody at 4□ overnight, followed by a horseradish peroxidase (HRP)-conjugated secondary antibody (1:5000). Protein bands were visualized using an SuperFemto ECL Chemiluminescence Kit (Vazyme, # E423-01). Images were captured by ChemiDoc MP (Bio-rad, RRID:SCR_019037) and analyzed using ImageJ software. Human α-actin and β-tubulin were used as a loading control. The antibodies and dilutions were listed in Suppl. Table S7.

### ELISA to detect the IL-8 concentration

After completing the co-culture experiments, culture supernatants were collected and stored at −80 °C for future analysis. The assays were conducted according to the manufacturers’ protocols (ELabscience, #E-OSEL-H0014). In brief, IL-8 was captured by antibodies pre-coated on 96-well plates. Subsequently, biotinylated detection antibodies were applied to the wells, followed by the addition of a streptavidin-horseradish peroxidase (HRP) complex. The presence of the IL-8 was then visualized by adding a substrate solution, and the absorbance was measured at approximately 450 nm using a spectrophotometer (Promega, RRID: SCR_018614).

### Whole transcriptome sequencing and data analysis

For RNA sequencing, total RNA was extracted from tumor cells using column-based purification method (Zymo, #R2052) and analyzed for quality on an Agilent 2100 Bioanalyzer. Ribosomal RNA (rRNA) was depleted using a commercially available kit (Vazyme, #N406-01), which selectively removes rRNA and enriches the mRNA fraction. The rRNA-depleted RNA was fragmented and used to construct a cDNA library following a protocol provided by VAHTS Universal V8 RNA-seq Library Prep Kit for MGI (Vazyme, #NRM605-01). The cDNA library was amplified using PCR and analyzed for size distribution on an Agilent 4200 TapeStation System (RRID:SCR_018435), then subjected to a circularization process to form a concatemers of the cDNA fragments (Vazyme, #NM201-01). After quantified using a Qubit fluorometer, the circularized library was loaded onto a sequencing flow cell and sequenced on a MGISEQ-2000 sequencer (MGI) in paired-end 150 bp mode.

RNA-seq data analysis was analyzed as in our previous study^62^. Briefly, raw sequencing reads were first subjected to adapter trimming and low-quality base removal with Ktrim software (version 1.5.0)^63^, and then aligned to the human reference genome (NCBI GRCh38) with STAR software (version 2.7.9a)^64^. Gene expression quantifications were conducted using FeatureCounts software (version 2.0.3)^65^. Subsequently, differential gene expression analysis was performed with DESeq2 (version 1.26.0)^66^. Gene Ontology (GO) enrichment analysis was conducted with ClusterProfiler (version 4.12.6)^67^. Gene Set Enrichment Analysis (GSEA) was applied to identify coordinated expression changes within biological pathways^68^.

### Immunofluorescence staining

For cell immunofluorescence (IF) staining, HCT116 cells were evenly distributed onto 8-well chamber slides at a density of 10^5^ cells per well. After reaching ∼90% confluence, cells were washed with phosphate-buffered saline (PBS) and 200uL serum-free medium supplemented with the 10^5^ *C. albicans* (MOI 1:1) was added. After 3 hours of culture in a humidified incubator (37°C/5% CO_2_), slides were fixed with 4% paraformaldehyde for 15 minutes at room temperature, followed by permeabilized with 0.1% Triton X-100 in PBS for 10 minutes. Non-specific binding was blocked with 5% bovine serum albumin (BSA) in PBS for 1 hour at room temperature. Cells were incubated with the primary antibody overnight at 4°C in a humidified chamber. After washing with PBS, the slides were incubated with a fluorochrome-conjugated secondary antibody (1:1000 dilution) and for 1 hour at room temperature in the dark. *C. albicans* were marked with Concanavalin A-Alexa Fluor 647 or Calcofluor White. Finally, the slides were mounted using an antifade mounting medium with DAPI (Beyotime, #P0131).

For organoids staining, uninfected and infected organoids were fixed in 4% paraformaldehyde in 100 mM PBS for 1 hour, washed with PBS with 100 mM glycine, permeabilized in 0.5% Triton X-100 in PBS for 1 hour, then incubated in staining buffer (4% BSA, 0.05% Tween-20 in PBS pH 7.4, 10% goat/donkey serum) for 4 hours, followed by incubation with primary antibody for 24 hours at 4°C. After washing with PBST (PBS with 0.05% Tween-20), incubated with fluorescent secondary antibodies, phalloidin and DAPI, for 4 h at room temperature. Following additional washes, whole mounts were submerged in antifade mounting medium (Beyotime, #P0126). Images were captured using Zeiss LSM900 confocal microscope (Carl Zeiss, RRID: SCR_022263) fitted with a 63× oil immersion objective or 10× objective.

### Construction of stable cell lines

Lentiviral particles were produced by co-transfecting 293T cells with the shRNA expression vector (pLKO-shRNA-EGFP vector) along with packaging (psPAX2) and envelope (pMD2.G) plasmids, using Lipofectamine 3000 Transfection Reagent (Thermos, #L3000015). The culture medium was replaced 24 hours post-transfection, and the lentiviral supernatant was collected 48 and 72 hours, filtered through a 0.45 µm filter. After HCT116 cells reach approximately 70% confluence, lentiviral particles were added to the cells in the presence of polybrene (5 µg/mL) to enhance infection efficiency. The cells were then incubated at 37°C for 24 hours. Stably transfected cells were selected using a Beckman CytoFLEX SRT Benchtop Cell Sorter (RRID: SCR_025068). The efficiency of shRNA-mediated knockdown was validated at the mRNA levels. The DNA primers for plasmid construction were listed in Table S1.

### Generation of C. albicans mutant strains

ECE deletion was performed according to the method descripted by Nguyen. Briefly, Guide RNAs (gRNAs) were designed using the CRISPOR online website (http://crispor.tefor.net/). The gRNA sequences were integrated with gRNA expression cassettes by cloning-free stitching PCR. The Cas9 expression cassette was amplified from vector pADH137 (RRID: Addgene_90986). The 500bp upstream and downstream sequences of ECE1 were fused with mCherry by fusion PCR. Cas9 cassette, gRNA cassette and donor DNA were transformed into the competent cells of *C. albicans* using the lithium acetate transformation method. Transformants were selected on YPD agar plates containing nourseothricin to isolate stably integrated cells. Single colonies from the selection plates were picked and grown in liquid YPD medium. Genomic DNA was extracted and the ECE1 locus modification was PCR amplified using primers flanking and inner of the target site.

### Tissue processing and organoid culture

Adjacent normal and tumor tissues were isolated from the resected colon segments. Healthy crypts and tumor epithelium was isolated essentially as described by Sato et al^69^. Organoids from adjacent colonic tissues were cultured in optimized human intestinal organoid medium^70^. The composition of medium is: advanced DMEM/F12 basal culture medium with 1% HEPES, 1% GlutaMAX, 1% Penicillin-Streptomycin, 10 ng/mL Wnt, 100 ng/mL R-Spondin, 100 ng/mL Noggin, 1× B27, 1,25 mM n-Acetyl Cysteine, 50 ng/mL human EGF, 5 μM A83-01, 3 μM SB202190, and 100 μg/mL Primocin. Tumor organoids were cultured with organoid medium minus Wnt and R-Spondin for the selection of organoids with intrinsic Wnt signaling activation. Organoids were split once a week and medium were refreshed every three days.

### H&E and immunohistochemistry staining

Tissue specimens collected from colorectal cancer patients were fixed in freshly prepared 4% paraformaldehyde at 4 °C for 24 hours. After fixation, the tissues were dehydrated, embedded, and sectioned. Hematoxylin and eosin (H&E) staining and immunohistochemistry (IHC) were performed using standard procedures. H&E staining involved the use of hematoxylin and eosin dyes. For immunohistochemistry (IHC), specific primary antibodies targeting β-1,3-glucan (1:100) and HIF-1α (1:200) were applied to detect the presence and localization of these markers within the tissue sections. The stained slides were scanned using the SLIDEVIEW VS200 system (Olympus, RRID: SCR_024783), and images were processed using OlyVIA software (Olympus, RRID: SCR_016167).

### Survival analysis

The abundance of *C. albicans* in colon adenocarcinoma samples in The Cancer Genome Atlas (TCGA) was collected from Narunsky-Haziza et al. study^71^, which characterized fungal populations across multiple cancer types^71^. Survival analysis was performed in the “survival” package in R. All experiments were performed in triplicate, and the data is presented as the mean ± standard error of the mean (SEM). Statistical analysis was conducted using unpaired t-test and a one-way ANOVA followed by a post-hoc Tukey test to determine the significance of differences between two groups and more than two groups respectively.

## Data availability

Raw RNA-seq data has been deposited to Gene Expression Omnibus (GEO) under accession number GSE279572.

## Acknowledgements

This work was supported by Noncommunicable Chronic Diseases-National Science and Technology Major Project (2026ZD0557400), Guangdong Basic and Applied Basic Research Foundation (2023B1515120073), National Key R&D Program of China (2022YFA0912700), China Postdoctoral Science Foundation (2023M742421), and Shenzhen Bay Scholar Fellowship (to C.D. and K.S.). We’d like to thank Ms. Qi Wang for technical assistance, SZBL Bioimaging Core, and SZBL High Performance Computing and Informatics Core for technical supports.

## Author contributions

Conception and design: W.W., K.S.; Study supervision: W.W., G.H., K.S.; Development of methodology: W.W., K.S.; Patient recruitment and clinical data analysis: C.D., N.L.; Cell culture and experiments: W.W., M.Y., Y.M.; Analysis and interpretation of data: all authors; Writing the manuscript: W.W., K.S.; Review and edit of the manuscript: W.W., M.Y., Y.Z., N.L., G.H., K.S.

