## Supplementary material for "*Candida albicans* drives colorectal cancer progression by inducing hypoxia signaling": Suppl. Figure

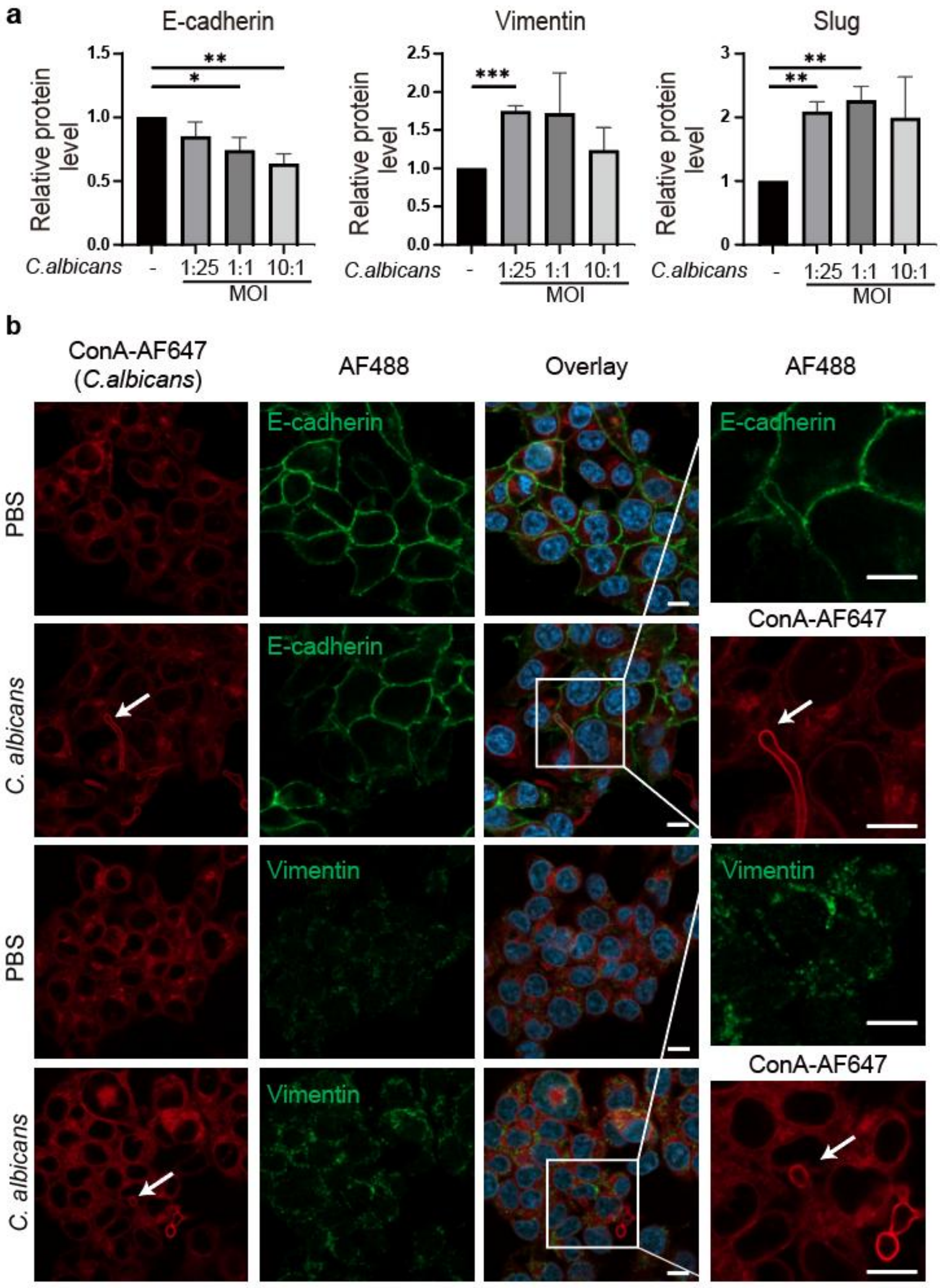


**Figure S1. *C. albicans* promote cell EMT. (a).** Statistical analysis of EMT protein level. Unpaired Student's t-test. *p < 0.05, **p< 0.01, ***p< 0.001. **(b).** Immunofluorescence images to detect EMT proteins of HCT116 cells. White arrowheads indicate hyphal of the C. albicans. White scale bar, 10 μm. Each treatment is a representative picture of at least three independent biological replicates.

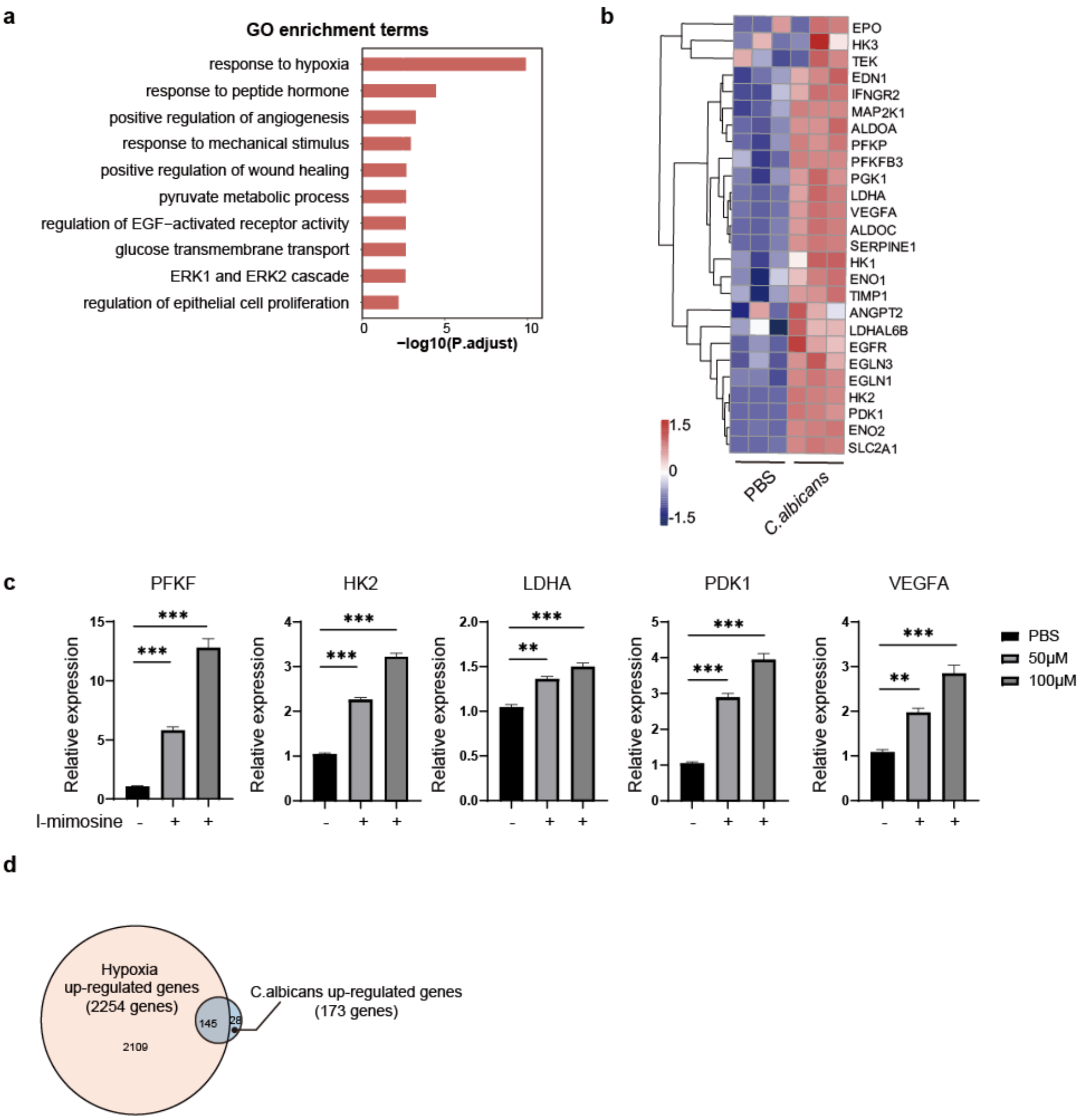


**Figure S2. C. albicans induces hypoxia response of HCT116. (a)** Gene Ontology enrichment analysis in HCT116 cells treated with C. albicans for 24 hours compared with PBS treatment. **(b)** Heatmap was drawn to show the HIF-1α downstream genes expression of HCT116 after C. albicans infection. **(c)** qRT-PCR was performed to detect gene expression of HCT116 cells treated with I-mimosine for 6 hours. **(d)** Overlap of differentially expressed genes between hypoxia condition and *C. albicans* infection in HCT116 cells. Unpaired Student's t-test. *p < 0.05, **p< 0.01, ***p< 0.001.

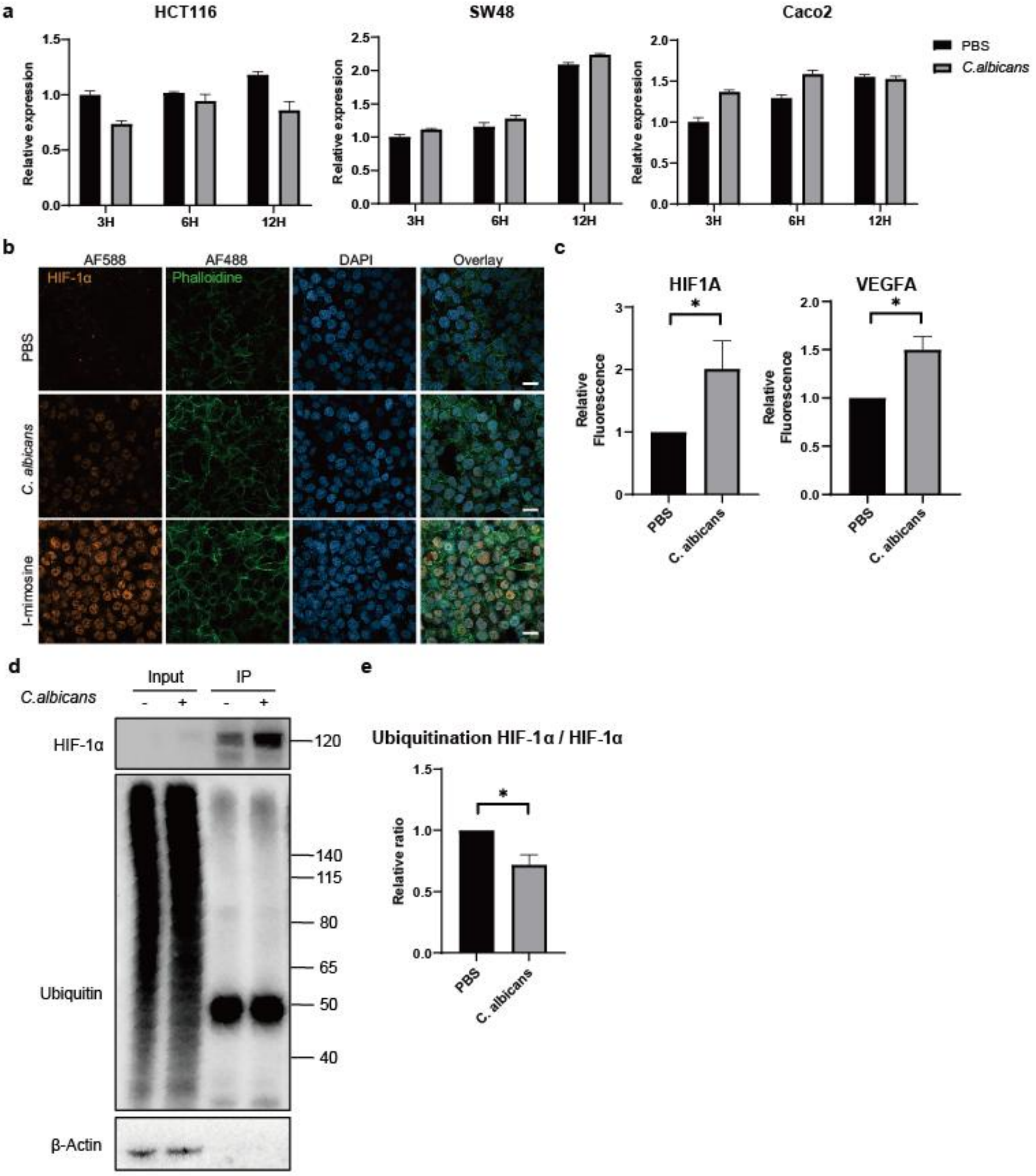


**Figure S3. C. albicans induces HIF-1α accumulation in HCT116 cells. (a)** qRT-PCR was performed to detect HIF1A expression of HCT116, SW48 and Caco2 cells infected by C. albicans after 3, 6, 12 hours. **(b)** Immunofluorescence images showing HIF-1α expression after *C. albicans* infection or i-mimosine treatment in HCT116 cells at 3 hours. The same scale bar was applied across all images in this panel. **(c)** Relative fluorescence level of HIF-1α and VEGFA of figure 2d, figure 2i. **(d)** Total and ubiquitinated HIF-1α levels following *C. albicans* infection were detected by immunoprecipitation. **(e)** High-molecular-weight bands (>50 kDa) in the IP lanes were quantified to represent ubiquitinated HIF-1α, whereas the ~120 kDa band was used to quantify total HIF-1α. The relative ratio of ubiquitinated HIF-1α to total HIF-1α was calculated. Unpaired Student's t-test. *$p < 0.05$.

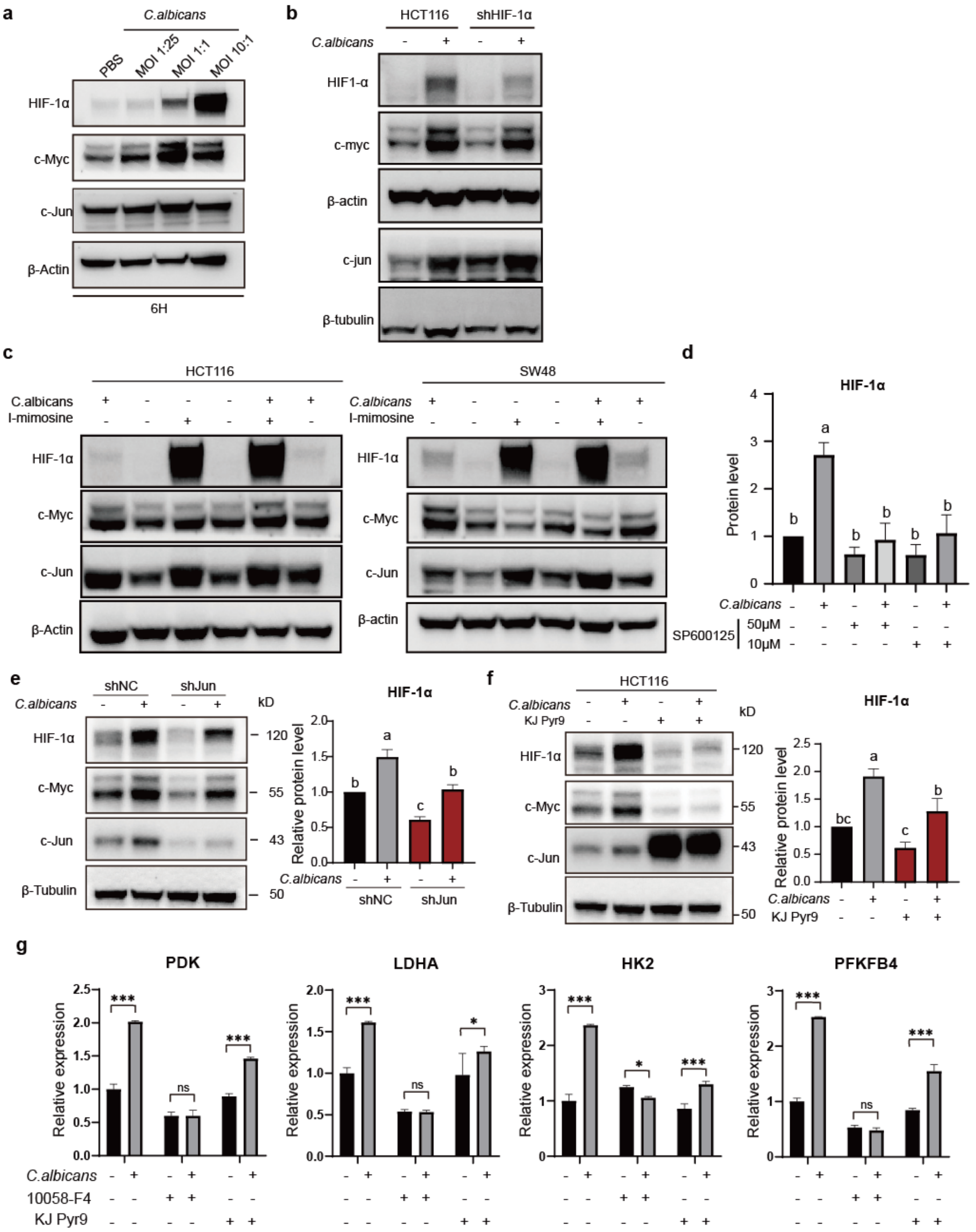

**Figure S4. c-Myc and c-Jun upregulation indispensable of HIF-1α. (a)** Western blot to detect HIF-1α, c-Myc and c-Jun of HCT116 infected by C. albicans with different multiple of infection (MOI). **(b)** Western blot to detect HIF-1α, c-Myc and c-Jun of wild type HCT116 and HIF-1α knock down cell lines infected by C. albicans. **(c)** Western blot to detect HIF-1α, c-Myc and c-Jun of HCT116 treated with i-mimosine or infected by C. albicans. **(d)** Statistical analysis of protein level of HIF-1α in HCT116 cells treated with c-Jun inhibitor SP600125. The results shown here are expressed as the mean ± SEM. Different letters indicate significantly different levels ($P < 0.05$) based on one-way ANOVA followed by Tukey's multiple comparison test. **(e)** Protein-level expression of HIF-1α, c-Myc, and c-Jun in c-Jun knock down cells after *C. albicans*

infection for 12 hours. **(f)** Protein-level expressions of HIF-1α, c-Myc, and c-Jun in HCT116 cells after *C. albicans* infection and treated with c-Myc inhibitor KJ Pyr9 for 12 hours. **(g)** qRT-PCR was performed to detect gene expression of HCT116 cells treated with *C. albicans* and c-Myc inhibitors for 24h. Two-way ANOVA followed by Bonferroni's multiple comparisons test. $*p < 0.05$, $**p < 0.01$, $***p < 0.001$.

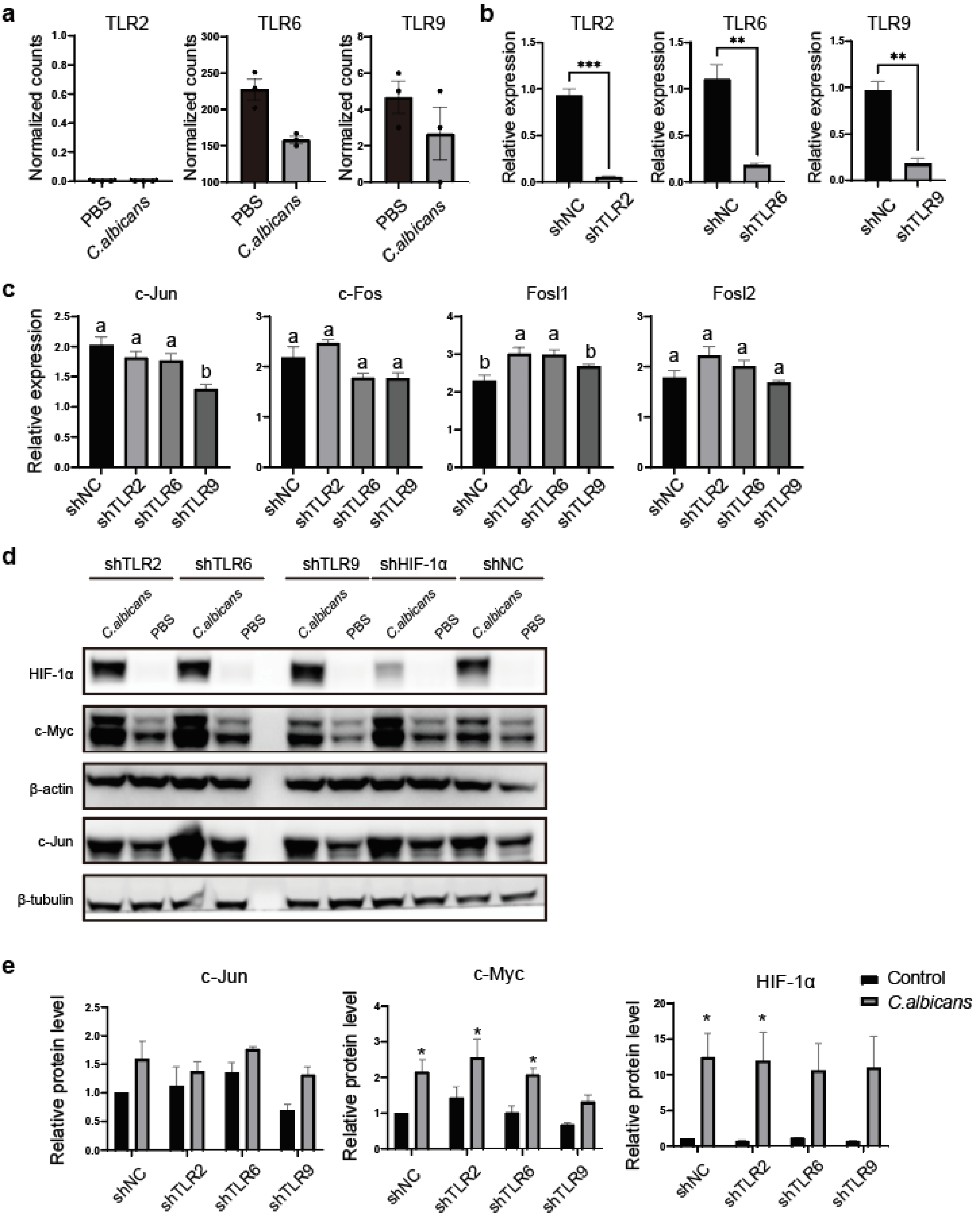


**Figure S5. TLR2, TLR6 and TLR9 gene knock down cannot inhibit the cell response triggered by C. albicans. (a)** Expression level of TLR with and without *C. albicans* infection. **(b)** qRT-PCR was performed to detect TLR knock down efficiency in different cell lines. Unpaired Student's t-test. $^*p < 0.05$, $^{**}p< 0.01$, $^{***}p< 0.001$. **(c)** qRT-PCR and western blot **(d)** was performed to detect gene expression in different cell lines infected with *C. albicans.* **(e)** Statistical analysis of protein level of different cell lines cultured with C. albicans for 12h. The results shown here are expressed as the mean ± SEM. ANOVA followed by Tukey's multiple comparison test. $^*p < 0.05$

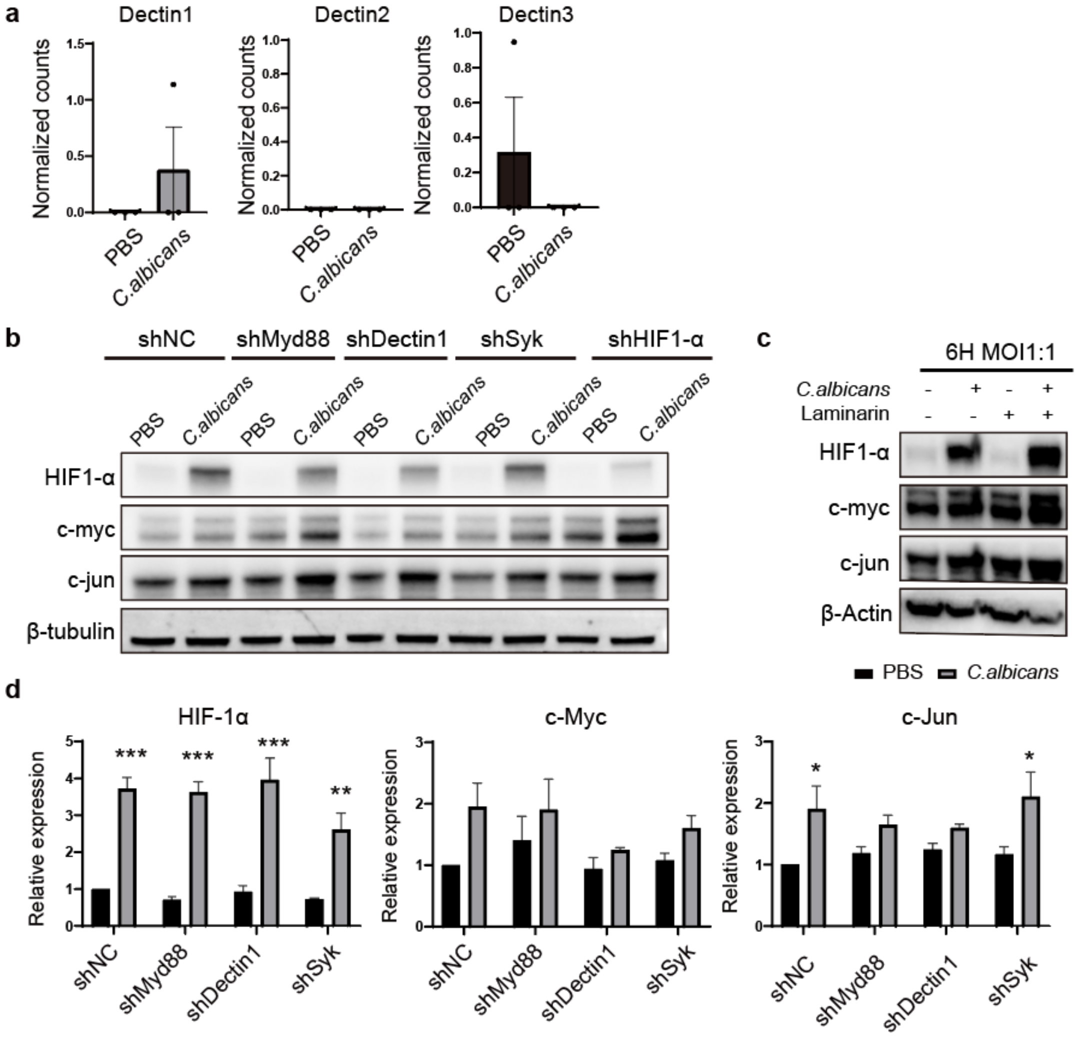


**Figure S6. CLR and adaptor proteins cannot inhibit the cell response triggered by C. albicans. (a)** Expression level of CLR with and without *C. albicans* infection. Unpaired Student's t-test. *p < 0.05.**(b)** Western blot was performed to detect gene expression in different cell lines infected with *C. albicans.* **(c)** Western blot was performed to detect gene expression in HCT116 cells infected with *C. albicans* with or without Laminarin treatment. **(d)** Statistical analysis of protein level of different cell lines cultured with C. albicans for 12h. The results shown here are expressed as the mean ± SEM. ANOVA followed by Tukey's multiple comparison test. *p < 0.05, **p< 0.01, ***p< 0.001.

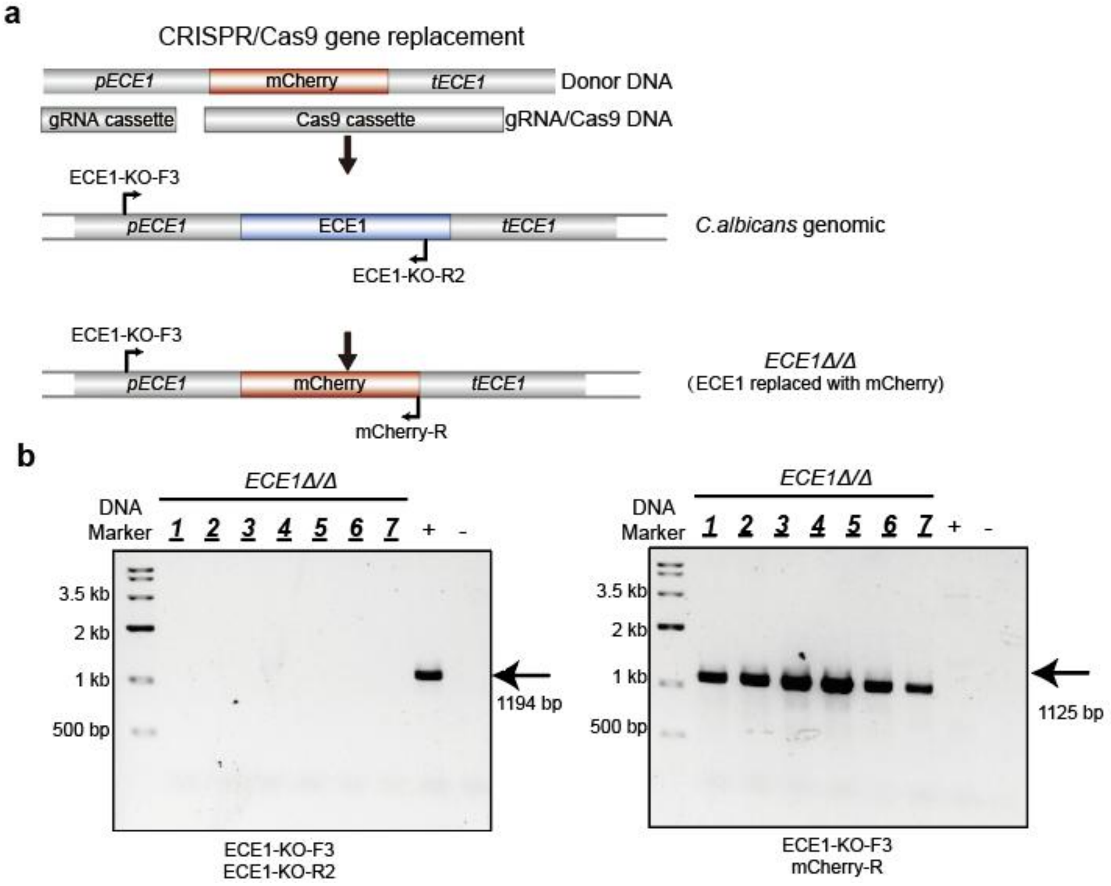


**Figure S7. ECE1 deletion strains construction and verification. (a)** Schematic diagram of knock out strain construction. **(b)** Gels show the verification DNA bands for the wild type (left panel) and mutant (right panel) amplified with indicated primers in **(a).** +: wild type strain, -: negative control (distilled water)

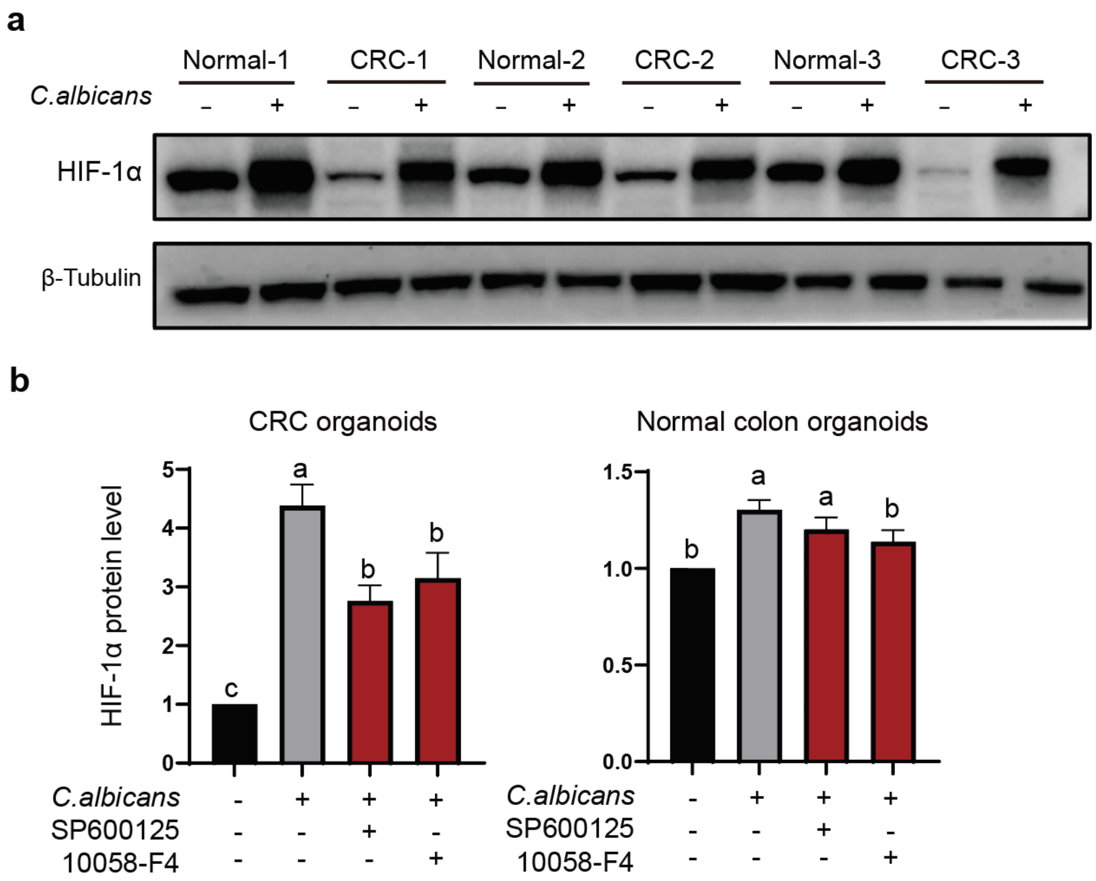


**Figure S8. *Candida albicans* induces hypoxia response pathway of normal colon organoids and CRC organoids. (a)** Western blot to detect HIF-1α proteins of normal and CRC organoids after C. albicans infection. **(b)** Statistical analysis of protein level corresponding to the organoids treated with c-Jun inhibitor SP600125 or c-myc inhibitor 10058-F4 and infected with *C. albicans* The results shown here are expressed as the mean ± SEM. Different letters indicate significantly different levels ($P < 0.05$) based on one-way ANOVA followed by Tukey's multiple comparison test.

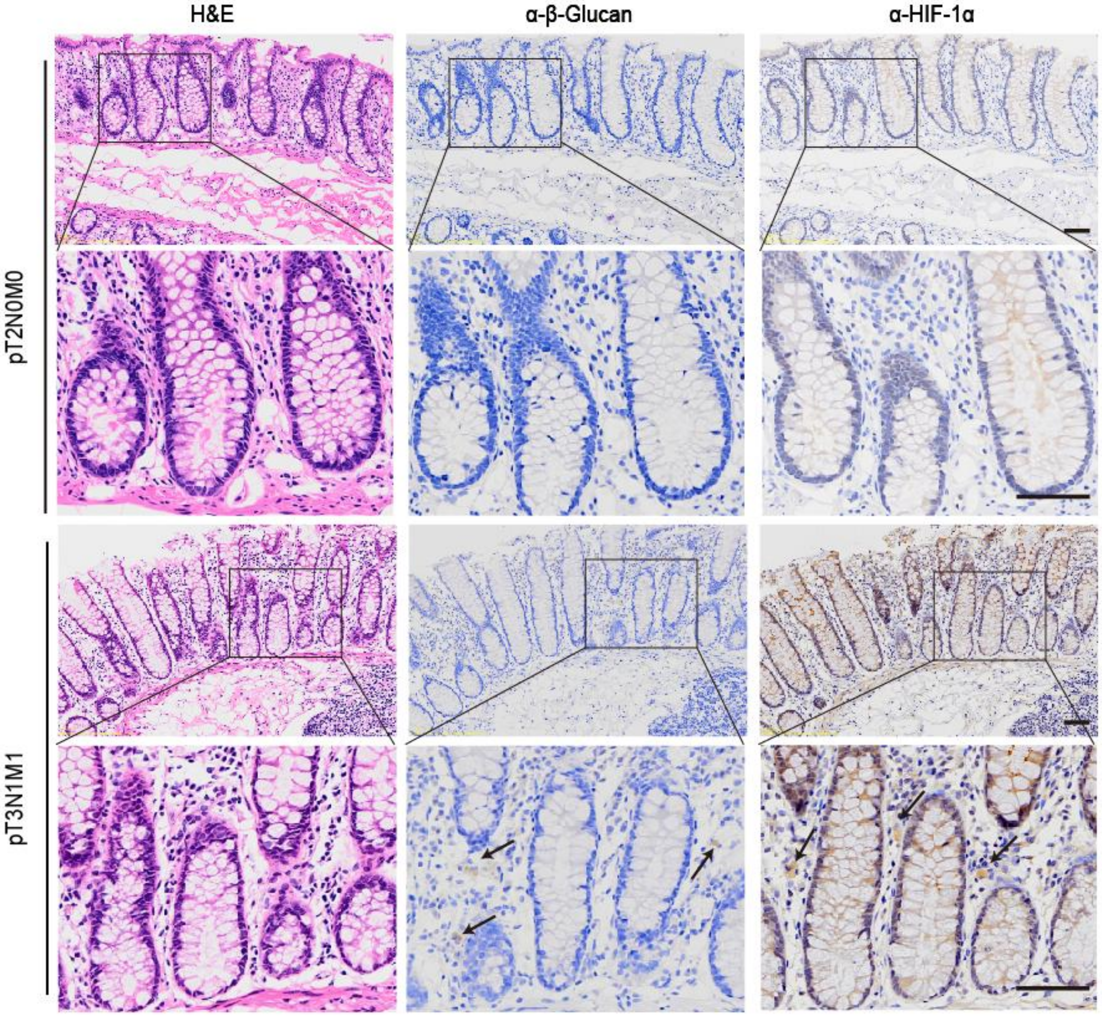


**Figure S9.** ***Candida albicans*** **induces hypoxia response pathway of CRC tissues.** Scale bars indicated 50 μm.

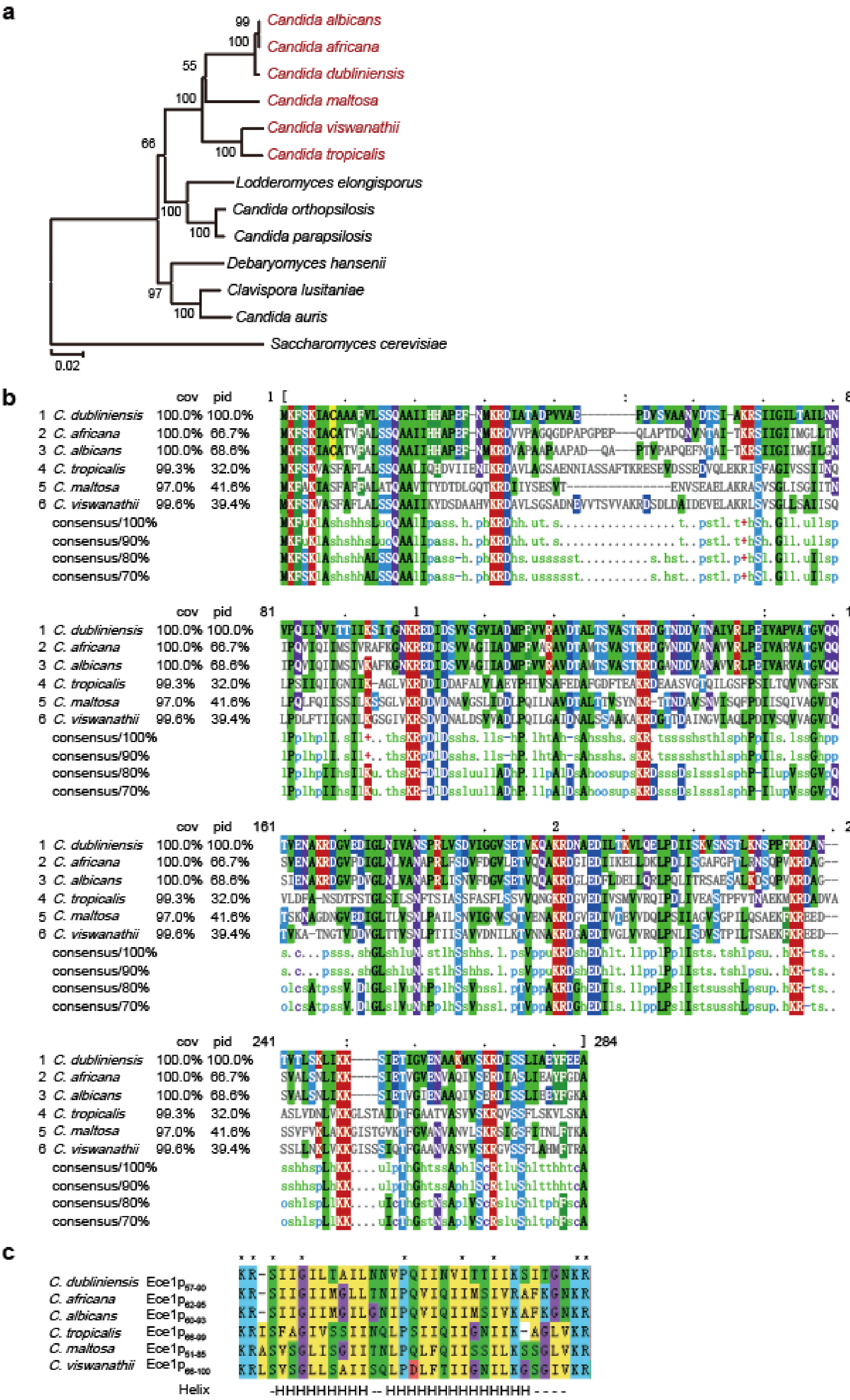


**Figure S10. Phylogenetic analysis of (a) ECE homolog, (b) multiple-sequence alignment of Ece1p, and (c) candidalysin-like peptides.**

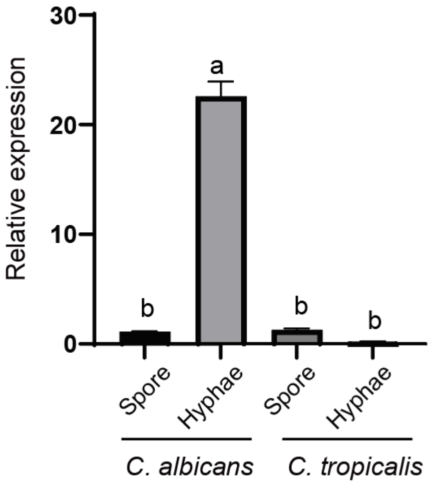


**Figure S11. ECE1 gene expression in C. albicans and C. tropicalis.** ECE1 gene expression in C. albicans, and C. tropicalis was quantified in the presence of HCT116 cells by RT-qPCR after 0.5 h.

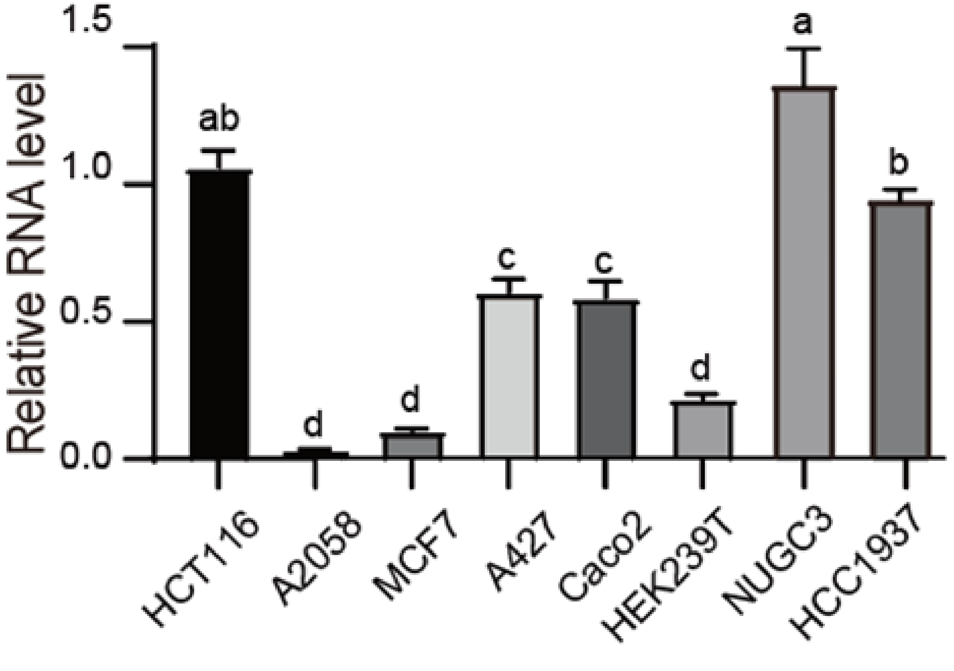


**Figure S12. Expression of EGFR in different cell lines.** The results shown here are expressed as the mean ± SEM. Different letters indicate significantly different levels ($P < 0.05$) based on one-way ANOVA followed by Tukey's multiple comparison test.
